# Genetic exploration of a nuclear receptor transcriptional regulatory complex

**DOI:** 10.1101/2020.03.28.013060

**Authors:** Masako Asahina, Deborah Thurtle-Schmidt, Keith R. Yamamoto

**Author notes:** Department of Biology, Davidson College, Davidson, NC, 28035. **Corresponding author:** Keith R. Yamamoto, 600 16th Street, University of California, San Francisco, GH-S572D, Box 2280, San Francisco, CA 94143-2280.

## Abstract

Metazoan transcriptional regulatory factors (TFs) bind to genomic response elements and assemble with co-regulators into transcriptional regulatory complexes (TRCs) whose composition, structure and activities are gene-, cell- and physiological-context specific. Each TRC is a “regulatory logic module,” integrating incoming signaling information, which defines context and thereby recruits a distinct combination of co-regulators that together specify outgoing regulatory activity. Analyzing TRCs unique to every context is daunting, yet justified by their properties as self-contained regulatory modules. As proof-of-concept, we performed a forward genetic screen in *C. elegans* carrying a synthetic simple response element for nuclear receptor NHR-25 upstream of a fluorescent reporter gene. We isolated independent mutations in *uba-2*, a component of the sumoylation signaling machinery, and in *lir-2*, which we demonstrated to be a novel co-regulator, interacting with NHR-25 through LxxLL motifs and modulating target gene expression. Our studies establish that an unbiased genetic screen readily identifies both afferent and efferent components that specify TRC function, and suggest that screening natural response elements of interest could illuminate molecular mechanisms of both context-specificity and transcriptional regulation.

## INTRODUCTION

Nuclear receptors (NRs) are transcriptional regulatory factors (TFs) that bind to genomic response elements and govern the expression of nearby genes essential for cell differentiation, development, and homeostasis (Yamamoto 1985; Kastner *et al.* 1995; Mangelsdorf *et al.* 1995). In metazoans, such complex phenotypes require many distinct transcription programs executed separately in different cells, tissues, organs and organ systems, and coordinated into functional networks. A key feature of that coordination is the capacity of a single TF, such as a NR, to specify gene expression programs that differ dramatically in different cell or physiological contexts (Weikum *et al.* 2017). The precision of each context-specific roster of NR-regulated genes is achieved by combinatorial assembly, at particular response elements, of transcriptional regulatory complexes (TRCs) comprised of a NR together with unique combinations of other TFs and coregulatory factors. In turn, the plasticity with which the roster of regulated genes changes with each new context is determined by signaling pathways that modify TRCs, and thereby their composition, structure, and activities.

NRs are modular proteins bearing a zinc-finger DNA-binding domain (DBD) coupled through a hinge region to a carboxy-terminal ligand binding domain (LBD). In addition to the non-covalent association with specific small molecules in the LBD (natural ligands have not yet been identified for all NRs), multiple positions on NRs are targets for a range of covalent post-translational modifications (PTMs), *e.g.,* phosphorylation, methylation, acetylation, ubiquitylation and sumoylation (Berrabah *et al.* 2011; Treuter and Venteclef 2011; Lee and Lee 2012). Ligands, PTMs, and even the precise DNA sequence with which an NR associates (Watson *et al.* 2013), all appear to serve as allosteric effectors, working together to alter NR conformation. Finally, multiple “receptor surfaces,” generated or stabilized by the allosteric effectors, confer transcriptional activation or repression; these are generally not well defined except for the C-terminal AF2 domain (Mangelsdorf *et al.* 1995). The regulatory surfaces function as docking sites for a broad range of co-regulatory factors (Lonard and O’Malley 2012), typically chromatin-remodeling ATPases (Yoshinaga *et al.* 1992) or enzymes that attach or remove small adducts (phosphoryl, methyl, acetyl, sumo) from TFs, other co-regulators, or neighboring nucleosomal histones; certain co-regulators may recruit other co-regulators or transcription initiation factors. Importantly, NR and other TFs have not been shown to carry intrinsic regulatory activities. Rather, TFs, including NRs, are DNA-binding scaffold proteins (Weikum *et al.* 2017) that acquire specific regulatory capacities by nucleating TRC assembly.

The characteristics of NRs and their interaction partners, together with the gene-, cell- and physiological-context specificity of their actions, identify NR-associated TRCs as “regulatory logic modules.” That is, these complexes are sole control points for afferent signals and efferent regulatory activities (Weikum *et al.* 2017), unlike the multiple control points that enable a membrane-bound receptor indirectly to modulate transcription in a conventional signal transduction pathway (*e.g.,* Budi *et al.* 2017). The realization that TRCs on the one hand are the molecular endpoints of upstream signals, commonly multiple signals, that specify their assembly, and on the other reveal molecular mechanisms of their regulatory activity, underscores the importance and impact of determining their structure and composition.

Two properties of TRCs limit the experimental and biological approaches to defining TRCs. First, they assemble on demand at context-determined response elements and exist only transiently; the turnover rate for a vertebrate NR TRC was shown to measure in seconds (McNally *et al.* 2000), underscoring the challenge of biochemical studies. Second, TRC composition is highly context dependent. Hence, an experimental system that enabled visualization of normal and aberrant TRC activity in different cellular, developmental and physiological contexts seemed essential. We chose, therefore, to pursue a genetic strategy in *C. elegans*. As proof of concept, we describe here a forward genetic screen utilizing a simple synthetic response element to identify signaling pathways and co-regulatory machinery that contribute to TRCs nucleated by NHR-25, a member of the highly conserved NR5A branch (also known as the Ftz-F1 family) of the NR superfamily, to which the human NRs SF1 and LRH-1 also belong (Antebi 2015). NHR-25 is broadly expressed throughout development and is involved in a range of biological functions such as cell fate determination and differentiation of somatic gonadal and epidermal cells, morphogenesis, molting, heterochrony, fat metabolism and stress responses (Gissendanner and Sluder 2000; Asahina *et al.* 2000; Chen *et al.* 2004; Silhankova *et al.* 2005; Asahina *et al.* 2006; Mullaney *et al.* 2010; Ward *et al.* 2013; 2014).

## MATERIALS AND METHODS

### C. elegans culture and strains

All *C. elegans* strains were cultured on normal nematode growth media (NGM) plates seeded with OP50 bacteria (Brenner 1974). The N2 Bristol strain was used as the wild type strain. The following strains were generated; HL178 *jmIs167[8NR5RE::pes-10∆::NLS::3xVenus::unc-54 3’utr + myo-2p::tdTomato]* (Ward *et al.* 2013), cr-155 is *HL218 lir-2 (jm218; jmIs167)*, cr-158 is HL219 *lir-2(jm219; jmIs167)*, cr-160 is HL220 *lir-2 (jm220; jmIs167)*, cr-169 is HL221 *lir-2 (jm221; jmIs167)*, cr-172 is HL222 *lir-2 (jm222; jmIs167)*, cr-178 is HL223 *lir-2 (jm223; jmIs167)*, cr-179 is HL224 *lir-2 (jm224; jmIs167)*. RW11401 strain was obtained from Caenorhabditis Genetics Center (CGC).

### Mutant screening

EMS (ethyl methanesulfonate, SIGMA-Aldrich) mutagenesis was performed essentially as described (Brenner 1974). Synchronized L4-stage worms of HL178 strain were treated with 50 mM EMS at 20 °C for 4 hr. Mutagenized P0 animals (about 20 worms per plate) were placed onto OP50-seeded NGM plates to further grow. Gravid P0 adults of each plate were sodium hypochlorite treated to collect eggs and F1 progeny was allow to grow on fresh OP50-seeded NGM plates. In the course of development, individuals with enhanced Venus expression were singled and screened further for the homozygosity in F2s. About 73,000 P0 gametes were screened and two independent mutant strains, EMS2 and EMS9, were isolated. Genomic DNA of EMS2, EMS9 and HL178 strains were isolated (Quick DNA MiniPrep Kit, Zymo Research), sheared using a water sonication bath (Bioruptor, Diagenode) and fragmented DNA was ligated to index adapters and amplified (BioO NEXTflex qRNA-Seq Kit v2, Perkin Elmer). Indexed DNAs smaller than 500 bp were gel purified and subjected to next generation sequencing (HiSeq 4000, Illumina). Quality and quantity of fragmented DNA and the library for sequencing were analyzed by Bioanalyzer (Agilent High Sensitivity DNA kit). To identify variants from the whole genome sequencing, samples were mapped to the ce6 genome using BWA (Li and Durbin 2009). For each strain, variants were detected using SAMtools mpileup and BCFtools for variant calling (Li and Durbin 2009). SNPs unique to the mutants were identified through comparison of the mutant-ce6 variants to that of wildtype-ce6 variants (github.com/dthurtle-schmidt/unique-snp-identification). Unique variants were annotated using ANNOVAR to identify those SNPs in coding regions (Wang *et al.* 2010). Number of reads mapped for each sample is recorded in Table S2. Reads have been deposited in the National Center for Biotechnology Information (NCBI) Sequence Read Archive (SRA) at http://www.ncbi.nlm.nih.gov/sra under accession no. PRJNA613114.

### CRIPSR-Cas9 genome editing

The *lir-2* locus was edited using CRISPR-Cas9 system according to (Paix *et al.* 2015; Dokshin *et al.* 2018), and Integrated DNA Technologies (IDT). Briefly, CRISPR-Cas9 ribonucleoprotein (RNP) complexes were preassembled by mixing 110 ng/μL tracr RNA (IDT), 100 ng/μL of each guide RNAs (Alt-R crRNA, IDT) and 250 ng/μL Cas9-NLS purified protein (MacroLab, QB3 Berkeley) and incubating at 37 °C for 10 min. 110 ng/μL of each ssODN donor templates (IDT) and 50 ng/μL pRF4: co-injection marker *rol-6 (su1006)* expression plasmid (Kramer *et al.* 1990) were then added to the complex mixture. The mix was microinjected into the gonadal arms of HL178 worms (Mello and Fire 1995). Injected P0 hermaphrodites were singled and allowed to lay eggs. F1 rollers (Rols) from jackpot-brood plate(s) based on the number of Rol animals in a plate were singled. On the second day after the initiation of egg laying, F1s were frozen individually in worm lysis buffer (10 mM Tris-HCl, pH 8.2, 50 mM NaCl, 2.5 mM MgCl_2_, 0.45% NP-40, 0.05% polyethylene glycol 8000) with 1mg/mL Proteinase K (Ambion) and subjected to single-worm PCR for genotyping. Worms are freeze-thawed 3 cycles and incubated at 65 °C for 1.5 hours followed by 95 °C for 10 min. Mutations were screened by MiSeq next generation sequencing (Illumina). CRISPR-targeted regions were amplified with MiSeq-compatible gene-specific primers containing Read1 and Read2 adaptor sequences (IDT) using KOD DNA polymerase Hot start kit (TOYOBO/Millipore). The second PCR was performed with index adaptor primers compatible with Ilumina MiSeq (i5 and i7 primers, IDT) using Phusion DNA polymerase. 5 μL of each PCR reaction from the single worm lysate was pooled and purified by SPRI (Beckman Coutler) to remove unincorporated oligos. Purified DNA was checked by biolanalyzer (Agilent) and quantified by qPCR with KAPA Std DNA and primers (KAPA) using Sso Advanced Universal SYBR premix system (Bio-Rad). MiSeq sample library was prepared according to the manufacture’s instruction (MiSeq reagent kit v2 Nano, Illumina) and loaded onto MiSeq. Sample sheet generation and sequence analyses were performed as described (Ehmsen *et al.* 2019). Progeny from the positive F1 with desired mutations was recovered and singled to get the homozygous mutant. Isolated mutants were backcrossed with N2 wild type males.

### RNA interference (RNAi)

Feeding RNAi was performed essentially as described (Timmons *et al.* 2001). dsRNA was initially induced for 4 hr in liquid culture using 1mM IPTG before bacteria were concentrated and seeded on plates containing 50 μg/mL carbenicillin, 12.5 μg/mL tetracycline and 1mM IPTG. Bacteria carrying pPD129.36 without an insert were used for control RNAi. Sodium hypochlorite-treated eggs were placed on RNAi plates seeded with dsRNA-induced bacteria. RNAi clones were recovered from RNAi libraries (Kamath *et al.* 2003; Rual *et al.* 2004) and sequence-verified before use. RNAi clone for *nhr-25* is as described previously (Silhankova *et al.* 2005).

### Imaging

For microscope imaging, worms were anesthetized in drops of 50 mM sodium azide (SIGMA-Aldrich), mounted onto 2% agar pads and observed under differential interference contrast (DIC) optics or fluorescent illumination (AxioPlan 2, Zeiss). For Venus expression, images were taken in black and white camera (Hamamatsu, ORCA-ER) and pseudo colored by ImageJ (NIH).

### Cell culture and Co-immunoprecipitation

Mycoplasma-free human embryonic kidney cells (HEK293T, ATCC) were maintained in Dulbecco’s modified Eagle’s medium (DMEM) supplemented with 10% fetal bovine serum (FBS, Gemini Bio) and antibiotics. Transfections were performed with the TransIT 2020 (Mirus). Full length wild type NHR-25 was tagged with EGFP at its N-terminus using pEGFP-C2 plasmid vector (Clontech); wild-type and mutant LIR-2s were N-terminally FLAG-tagged using the p3xFLAG-CMV-10 expression vector (SIGMA-Aldrich). For co-immunoprecipitation, cells were harvested 36 hr after transfection, washed twice in ice-cold PBS and lysed in lysis buffer (50 mM Tris-HCl (pH 7.4), 150 mM NaCl, 1% Triton X-100, 1 mM EDTA and proteinase inhibitor (cOmplete, Roche) for 30 min at 4 °C on a rocking platform followed by centrifugation at 15,000 rpm for 15 min at 4 °C. The supernatant was transferred to a fresh tube and incubated with anti-GFP antibody coupled with magnetic agarose (GFP-Trap, Chromotek) overnight at 4 °C on a rocking platform. Beads were washed twice (50 mM Tris-HCl (pH 7.4), 150 mM NaCl containing proteinase inhibitor); proteins were eluted in SDS sample buffer with 10% 2-Mercaptoethanol (Bio-Rad), separated on SDS-PAGE, and transferred to Immun-Blot PVDF 0.2 μm membrane (Bio-Rad). FLAG-tagged LIR-2 protein was detected with monoclonal anti-FLAG M2-Peroxidase (HRP) antibody (SIGMA-Aldrich) and EGFP:NHR-25 was visualized with Clarity Western ECL substrate (Bio-Rad). Images were taken with Bio-Rad ChemiDoc Imaging system.

## DATA AVAILABILITY

Sequence reads are deposited in the National Center for Biotechnology Information (NCBI) Sequence Read Archive (SRA) at http://www.ncbi.nlm.nih.gov/sra under accession number PRJNA613114. Strains and plasmids are available upon request. Supplemental materials are available through the GSA figshare portal.

## RESULTS

### Forward genetic screen for factors affecting NHR-25 transcriptional regulatory activity

NHR-25’s transcriptional regulatory activities are influenced by numerous signaling systems, including the Wnt/β-catenin (Silhankova *et al.* 2005; Asahina *et al.* 2006; Hajduskova *et al.* 2009) and heterochronic pathways (Hada *et al.* 2010; Nelson *et al.* 2011; Monsalve and Frand 2012), microRNAs (Hayes *et al.* 2006), phosphoinositides (Mullaney *et al.* 2010), and posttranslational modifications such as sumoylation (Ward *et al.* 2013). We showed previously that NHR-25 binds to the SF-1/LRH-1 (NR5A) consensus DNA binding motif (NR5RE: (C/T)CAAGG(C/T)CA) *in vitro*, and that a fluorescent reporter construct carrying that element, 8xNR5RE(WT)::3xVenus (Fig. 1A), was activated in *C. elegans* epithelial cells upon RNAi-mediated reduction of *smo-1* (Ward *et al.* 2013), the SUMO homolog that modifies target proteins in the sumoylation process. Thus, this minimal response element, which binds only a single TF in *C. elegans*, is functional *in vivo* and sensitive to at least one signaling pathway. We therefore used it in a forward genetic screen, testing for Venus reporter expression under conditions in which NHR-25 activity is normally suppressed, to identify additional components of an NHR-25 TRC. To begin, we chromosomally integrated the reporter, facilitated by UV irradiation, and backcrossed five times with wild-type strain N2, to produce strain HL178.

**Fig. 1.**
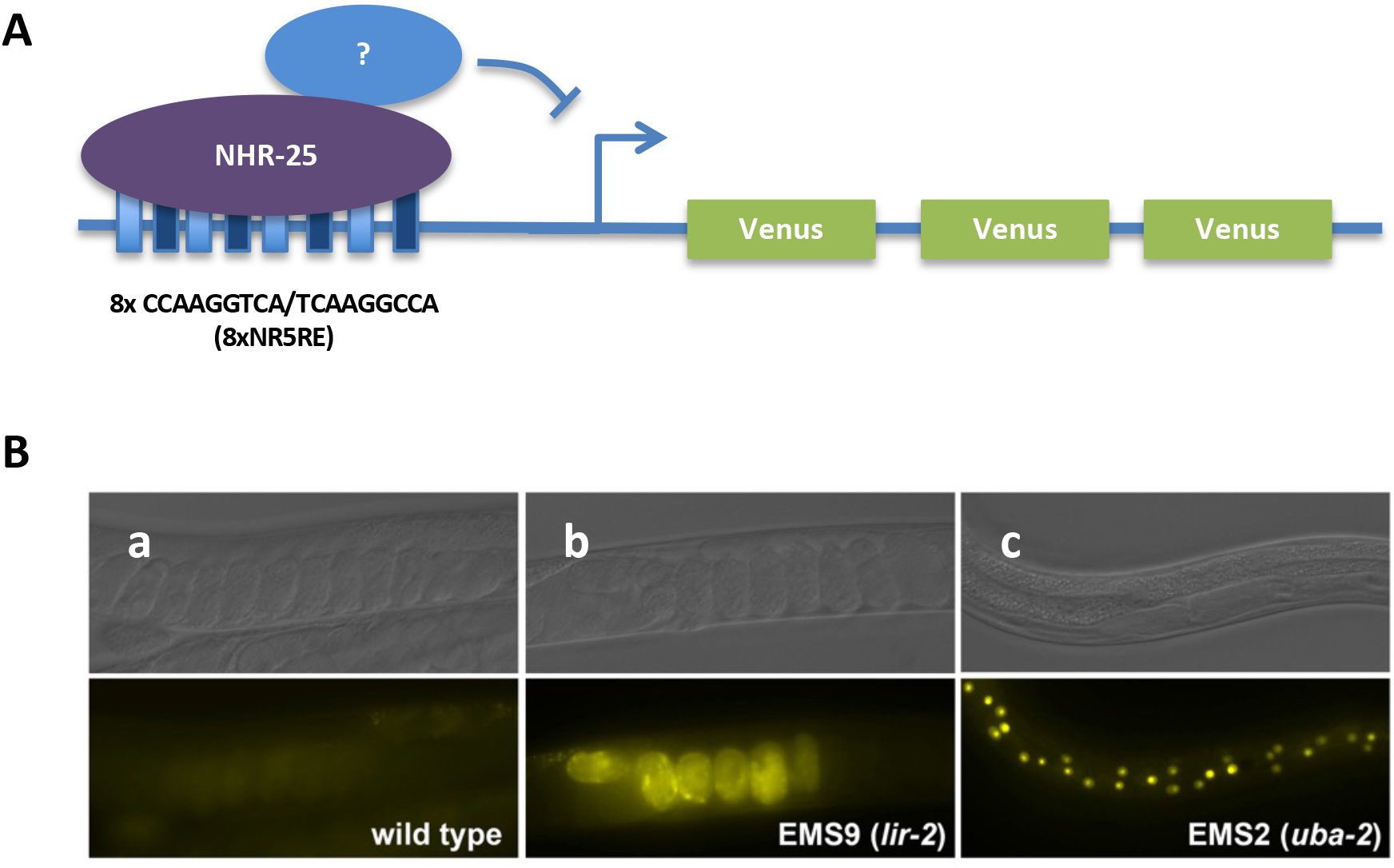
EMS forward genetic screen uncovered two independent genes regulating 8xNR5RE::Venus expression. (A) Schematic drawing of the transgene construct in HL178 strain, (B) 8xNR5RE::3xVenus transgene in wild type background shows very low level of expression in embryos (a), two independent mutant strains isolated in this screen (EMS9 and EMS2) show upregulated Venus expression in embryos (b) and larval epidermal cells (c). Deep sequencing revealed mutations in *lir-2* and *uba-2* genes, respectively.

Synchronized L4 HL178 hermaphrodites were ethyl methanesulfonate-mutagenized and Venus expressing individuals were isolated and singled. From 73,600 P0 gametes screened, we obtained two independent mutant strains, EMS2 and EMS9. In EMS2, Venus expression was observed in NHR-25-expressing (*e.g.,* lateral seam, hyp7 and vulval epithelial cells) (Fig. 1B-c). In EMS9, the reporter expression was seen both in embryos (Fig. 1B-b) and in larval epithelial cells. Through whole-genome sequencing of EMS2, EMS9, and HL178, we identified putative causative mutations for EMS2 and EMS9 in the *uba-2* and *lir-2* genes, respectively. To test if reduction in function by RNAi would phenocopy the identified mutants, we fed RNAi bacteria expressing dsRNA corresponding to *uba-2 or lir-2* sequences to HL178. These results showed that reduction of function in either of these genes produced activation of the Venus reporter.

### UBA-2 is a component of the sumoylation machinery that signals to NHR-25

*uba-2* encodes Ubiquitin-like modifier activating enzyme 2 (also known as SUMO-activating Enzyme subunit 2 or SAE2). UBA2 and its heterodimeric partner AOS1 (or SAE1) form an E1 activating enzyme, one of many components involved in sumoylation (Fig. 2A). The *C. elegans* genome encodes orthologs of each human component of the sumoylation machinery, and *C. elegans* UBA-2 is homologous to human UBA2 (42% identity, 57.4% similarity by EMBOSS Needle pairwise sequence alignment). The mutation in EMS2 substitutes an isoleucine with phenylalanine at amino acid 383, which resides between the Cys-domain and UbL-domain, a region highly conserved from yeast to human (Fig. 2B, C). The crystal structure of the E1 complex Sae1/Sae2 (Lois and Lima 2005) reveals that the region between the UbL and catalytic Cys domains is the physical interface with SUMO-1 protein and therefore essential for function of the E1 activation enzyme. As suggested by the RNAi results, it is likely that the EMS2 isolated mutation is a loss-of-function allele in *uba-2*.

**Fig. 2.**
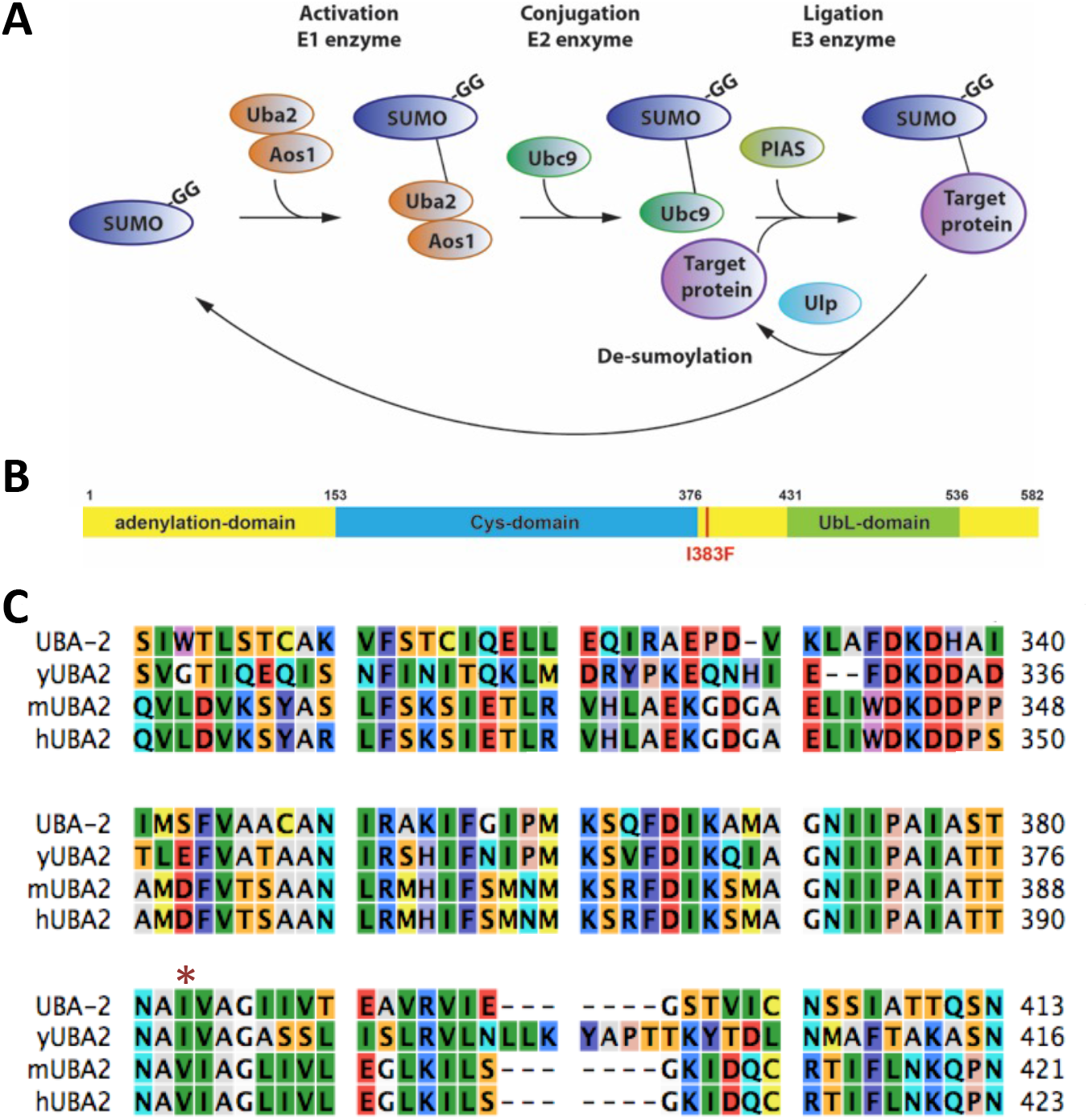
UBA-2 is a conserved component of sumoylation machinery. (A) Series of SUMO protein conjugation occur through Aos1/Uba2 E1 activation complex, E2 enzyme Ubc9 and E3 ligase PIAS to modify target protein. (B) Protein structure depicted based on the conservation with human UBA2. Mutation is located at the amino acid position 383 indicated in red (I383F). (C) Amino acid 315-413 of *C. elegans* UBA-2 was aligned with yeast, mouse and human UBA2 counterpart. Mutation (marked with asterisk) lays in the highly conserved region. Amino acid residues are colored with Rasmol colors based on protein properties.

Together with our previous report (Ward *et al.* 2013), we now know that two genes functional in sumoylation can modulate NHR-25-dependent transcription *in vivo*. This prompted us to test other components necessary for sumoylation, namely AOS-1 (E1 enzyme partner), UBC-9 (E2 conjugation enzyme), GEI-17 (E3 ligase), as well as the de-sumoylation Ulp family: ULP-1, ULP-2 and ULP-4 (Fig. 2A). Sodium hypochlorite-treated HL178 eggs were placed on feeding RNAi plates seeded with each corresponding bacterial clone. As expected, interference with sumoylation triggered robust upregulation of 8xNR5RE (WT)::3xVenus transgene expression in the seam cells and hyp7 cells (Fig. 3), whereas inhibition of de-sumoylation through RNAi of *ulp-1*, *ulp-2* and *ulp-4* had no effect, thus confirming that sumoylation inhibits NHR-25-mediated transcriptional activation of our 8XNR5RE reporter construct.

**Fig. 3.**
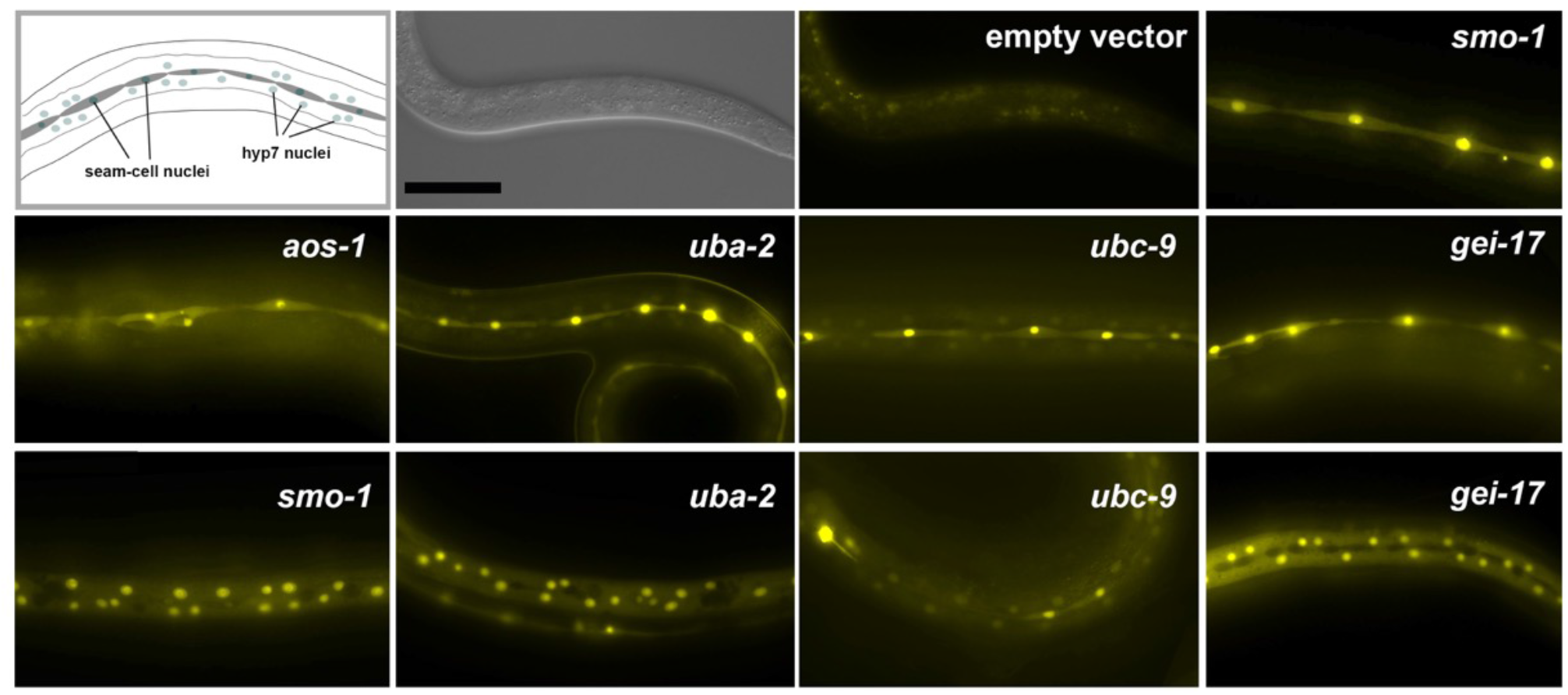
Components of SUMO conjugation machinery regulate NHR-25-dependent transcription. Left corner depicts the morphology of epidermal cells namely lateral seam cells (darker gray) and hyp7 cells (light gray nuclei). Targeted genes for RNAi are indicated in each panel. Upregulation of Venus expression seen in the seam cells (top panel of *smo-1* and all middle panels) and hyp7 cells (bottom panels) are shown. *uba-2* RNAi recapitulates the EMS mutant (EMS2) expression. Scale bar applies to all panels (50 μm).

### LIR-2 is a novel co-regulator of NHR-25

*lir-2* encodes <u>l</u>in-26-<u>r</u>elated Zn-finger protein (Dufourcq *et al.* 1999), and maps upstream of another *lin-26*-related gene *lir-1* and *lin-26* (Fig. S1). LIR-2 is highly conserved among Nematoda, but no clear homologs are found outside of that phylum. It contains a C2H2 Zn-finger motif, perhaps implying a DNA binding activity, as with LIN-26 (Fig. 4A). Unlike LIN-26, which is essential for epithelial cell differentiation and maintenance, and loss of function produces embryonic lethality, loss of function alleles of *lir-2* (*tm560*; National BioResource Project and EMS9; this study) is homozygous viable with no apparent morphological defects.

**FIG. 4.**
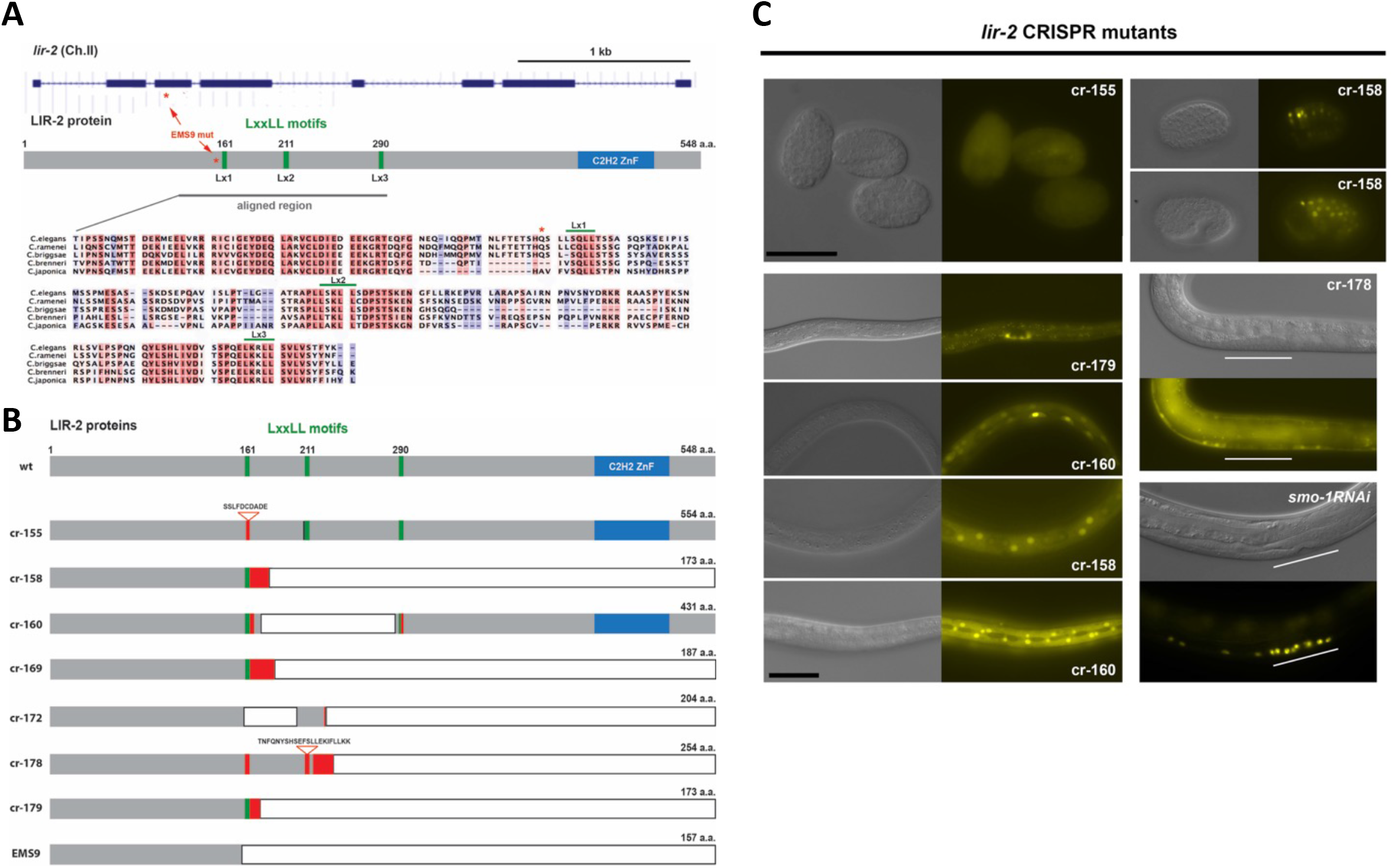
LIR-2 may be a coregulator of NHR-25. (A) *lir-2* genomic and protein structures are shown. Mutation S9 is indicated in red asterisk. Three LxxLL motifs are found at the position 161, 211 and 290 in LIR-2 and marked in green. The region containing LxxLL motifs (Lx1, Lx2 and Lx3) were aligned with ogous proteins in other Nematode species and showed that the region is highly conserved. LxxLL motifs icated over the amino acid sequences. (B) LIR-2 Protein structure of CRISPR-Cas9 generated mutants. dicates alteration in sequence, triangle indicates insertion, and deletion is marked as a white box. (C) sentative Venus expression in different tissues in mutants were shown. White bars in cr-178 and smo-1 panels indicate position of P5-7.pxx vulval cells. Scale bars: The black bar at the left top applies to all raphs of embryos and the black bar at the left bottom applies to all larval micrographs (50 μm).

In addition to its Zn-finger domain, LIR-2 carries three centrally-located LxxLL motifs (where x can be any amino acid), which are highly conserved in evolutionary distant nematode species such as *C. briggsae* and *C. japonica* (Fig. 4A). LxxLL is a highly conserved signature sequence that serves as the interaction surface on co-regulator factors for interaction with nuclear receptors; indeed, it is also denoted the NR-box (see Discussion). The EMS9 allele of *lir-2* is a C to T transition in the third exon that introduces a termination codon in place of Gln158, three amino acid residues upstream of the first LxxLL motif, generating a truncated protein that lacks all three LxxLL motifs (Fig. 4A and B).

To characterize further the role of *lir-2* in specifying NHR-25 regulatory activity, we created a series of Cas-9-mediated *lir-2* mutations. Although repair templates were injected (Table S1), efficiency of homology-directed repair was apparently low, as all recovered mutations were indels (cr-155 to cr-179; Fig. 4B) generated by non-homologous end joining: small insertions or deletions at the cleavage sites, as well as larger deletions (49 – 406 bp) close to the targeted PAM site at each locus (Fig. S2). Similar to EMS9, five of the lines, cr-158, cr-169, cr-172, cr-178 and cr-179, introduce termination codons that truncate the LIR-2 protein close to Lx1, the first LxxLL motif. The cr-155 line carries an in-frame 10 aa insertion that disrupts Lx1, and a deletion of a lysine residue immediately downstream of Lx2, which likely remains functional as another lysine follows the deletion (Figs. S2 and S3). The cr-160 line carries three mutations that together maintain the reading frame: a 358 bp deletion followed by a 7 bp insertion, which delete Lx2 and substitute four amino acids beginning six residues downstream of Lx1; in addition, Lx3 was mutated by HDR, LxxLL to LxxAA (Figs. S2 and S3). As amino acid residues flanking LxxLL are thought also to be important for NR recognition (McInerney *et al.* 1998), all three LxxLL motifs might be functionally compromised in this strain.

All of the alleles were homozygous viable, and all except cr-155 upregulated the NHR-25 reporter gene with indistinguishable expression patterns (representative examples in Fig. 4C): hypodermal cells in embryos; Z1/Z4 (and their daughters) somatic gonad precursors (SGPs), and P cells in early larvae; and lateral, dorsal, head and tail hypodermal cells throughout the larval stages. While all of these cells and tissues express NHR-25, NHR-25 expression alone was not sufficient for Venus upregulation. Thus, vulval precursor cells (P5-7.p and daughters) express NHR-25 robustly, but lack Venus expression (Fig. 4C, cr-178 panel). Similarly, NHR-25 expression is higher in seam cells than other hypodermal cells, yet seam cell Venus expression was only occasionally detected, and then only in dividing seam cell nuclei (Fig. 4C, cr-158 panel). Interestingly, strong reporter expression was seen both in the seam cells and vulval cells when SUMO machinery was compromised in the same strain (Fig. 3 and Fig. 4C, *smo-1* RNAi panel). This result confirms that NHR-25 is functional and signal responsive in each cell type, and that SUMO-dependent down regulation of NHR-25 is not LIR-2 dependent. Rather, the mechanisms by which SUMO and LIR-2 affect NHR-25 function are likely different.

Notably, Venus expression was weakly upregulated in cr-155 embryos (Fig. 4C, cr-155 panel), but no larval expression was detected. This implies that LIR-2 effects on NHR-25 activity depend more strongly on Lx1 in embryos than in the larval context. In general, it seems likely that each of the LxxLL motifs contribute to LIR-2 action on NHR-25, albeit to different extents depending on context.

To test if the upregulated Venus expression in *lir-2* mutants is NHR-25 dependent, we fed our *lir-2* mutant lines with bacteria expressing dsRNA sequences from *nhr-25*. Fig. 5 presents examples from cr-169 and cr-172: Empty vector control animals expressed the reporter in hypodermal cells (lateral and ventral hyp7 cells are shown), while expression was abrogated in *nhr-25* RNAi-treated animals. Thus, Venus activation in the *lir-2* mutants is NHR-25 dependent. We noted that expression in vulval cells (marked with asterisk) appeared to be unaffected, likely indicating that the high level of NHR-25 expression in those cells was not reduced sufficiently by RNAi.

**Fig. 5.**
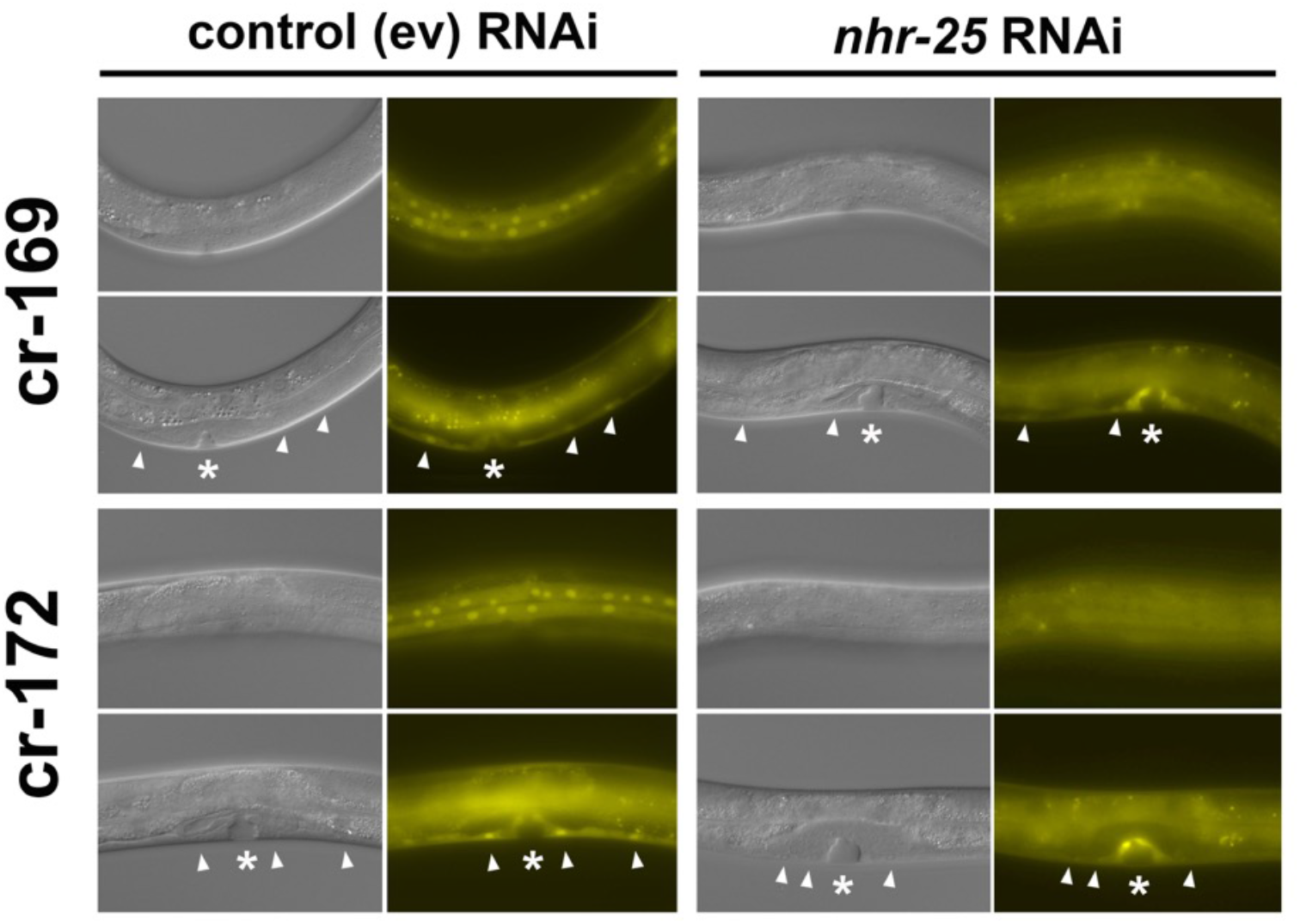
Upregulation of 8xNR5RE::Venus transgene expression is NHR-25 dependent. Left columns are control RNAi (empty vector or ev)-treated individuals showing the strong Venus expression in both lateral and ventral hyp7 nuclei in cr-169 and cr-172 *lir-2* mutants. Right panels are *nhr-25* RNAi-treated individuals showing the diminished Venus expression in hyp7 cell nuclei. Vulva position indicated with white asterisks and ventral hyp7 cells are marked with white arrowheads.

### LIR-2 physically interacts with NHR-25

Our genetic studies suggest that LIR-2 acts as an NHR-25 co-regulator, associating with the receptor through its LxxLL motifs. Accordingly, we tested their physical interaction by co-immunoprecipitation in HEK 293T cells. Full length of NHR-25 was tagged with EGFP and co-transfected with LIR-2 wild type, an E9 truncation that mimics the EMS9 variant (Figs. 4B, 6B), and an engineered mutant with LxxAA at Lx1, Lx2 and Lx3, each FLAG-tagged and expressed from the CMV10 vector (Fig. 6B). NHR-25 and presumptive protein complexes were precipitated with anti-GFP coupled to nanobody-conjugated magnetic beads, and LIR-2 protein was detected on western blots using anti-FLAG antibody conjugated with HRP. As shown in Fig. 6A, wild-type LIR-2 co-immunoprecipitates with NHR-25, while the E9 truncation does not, and the Lx1,2,3 mutant is drastically attenuated in binding capacity. We conclude that LIR-2 binds to NHR-25 and co-regulates its activity, functioning as a down-regulator at the 8xNR5RE response element in various cell- and development-contexts.

**Fig. 6.**
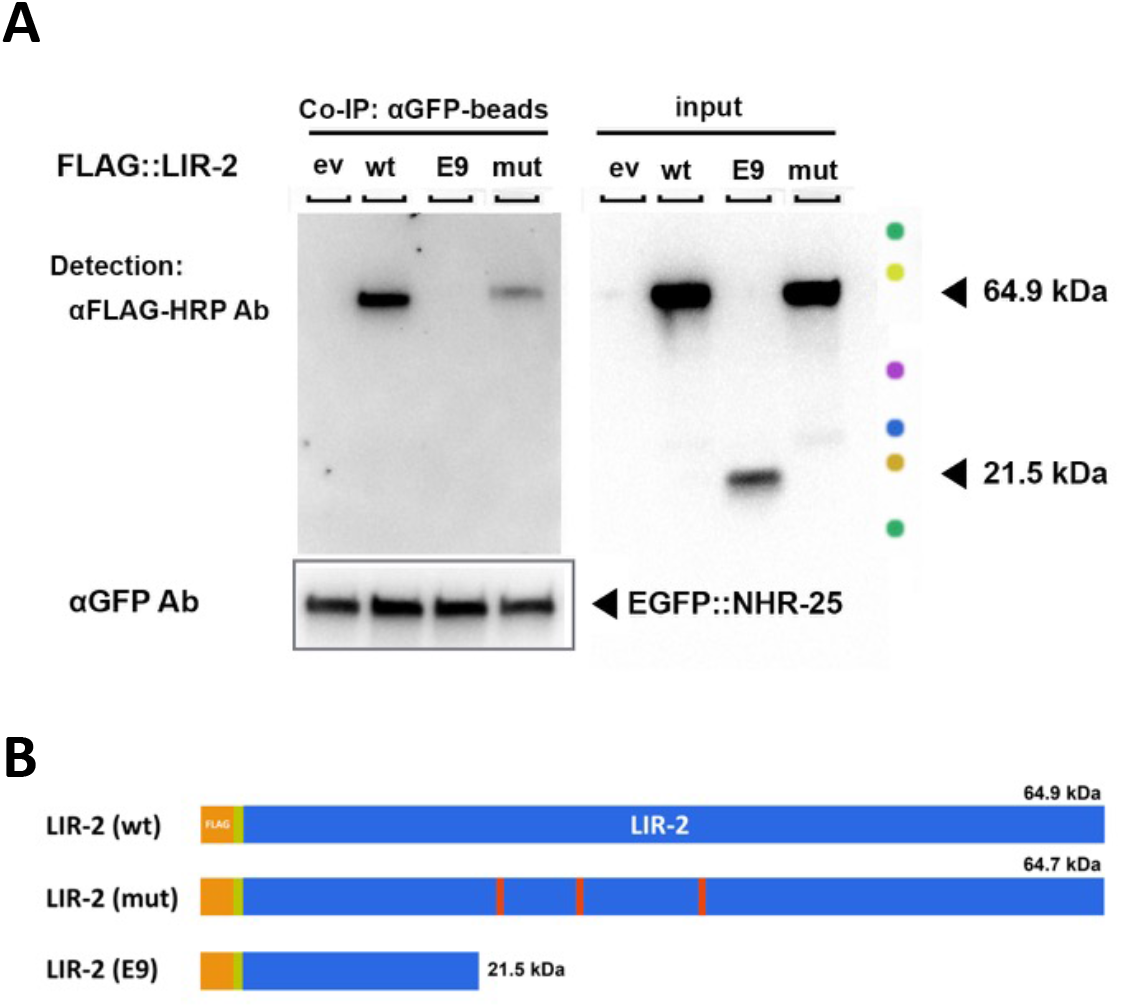
LIR-2 and NHR-25 physically interacts. (A) Co-immnopreciptation was performed in HEK293T cells. GFP-tagged NHR-25 and FLAG-tagged LIR-2 were co-expressed in the cells, precipitated with anti-GFP-nanobody conjugated agarose beads, and LIR-2 protein was detected by anti-FLAG antibody on western blotting. Wild type LIR-2 but not truncated protein (mimicking the EMS9 mutant protein) was bound to full length of NHR-25. Mutations in LxxLL motifs in LIR-2 protein remarkably diminished their physical interaction. (B) Schematic drawing of the LIR-2 protein variants tested in the co-immunoprecipitation assay.

## DISCUSSION

Metazoan transcription is governed by TFs that drive different patterns of expression in different cellular and physiological contexts, generating organismal networks of expression that give rise to complex biological processes, *e.g.,* development, metabolic homeostasis, responses to nutrients or stressors. Coordination of many distinct programs across tissues and organs to produce a coherent process requires that a given TF specify a precise expression pattern in each context, while still capable of great plasticity to confer drastically different patterns in different contexts. These seemingly contradictory properties of precision and plasticity are housed in context-specific transcriptional regulatory complexes that assemble transiently on-demand at genomic response elements (Weikum *et al.* 2017).

Importantly, a TRC is a regulatory logic module — concurrently the end point of multiple signal transduction pathways and the determinant of a regulatory mechanism emergent from combinatorial occupancy by specific subsets of co-regulatory factors. Thus, as the sole nexus of plasticity-enabling signaling and precision-specifying regulation, characterization of TRC components will illuminate both. The magnitude of such analyses is daunting, however, as TRCs likely comprise roughly 10^2^ polypeptides (Weikum *et al.* 2017) and their composition is influenced by multiple multi-step signaling pathways. In principle, a genetic screen for dysfunction of a response element would identify both TRC components and signaling machineries relevant to the context of the screen.

We report here a proof-of-concept, in which we sought mutations in genes that down-regulate a synthetic NHR-25 response element, 8xNR5RE, comprised solely of two NHR-25 consensus binding sequences in tandem: 4x CCAAGGTCA and 4x TCAAGGCCA (Ward *et al.* 2013). That simple element confers sumoylation-sensitive NHR-25-mediated regulation in a reporter context (Ward *et al.* 2013). We chose NHR-25 for this study because it is expressed in many cell types throughout development, and controls temporally a wide range of complex actions (*e.g.* cell fate decision and differentiation, fat metabolism, neuron projection and dauer formation). Finally, *C. elegans* was our metazoan of choice because it permits visualization of TF activity at specific cell resolution in embryos, larvae and adults across the full lifespan.

Our screen yielded both a signaling machinery component and a novel co-regulator, both of which down-regulate NHR-25 dependent expression in the context of the screen. Mutation of *uba-2* gene, a component of the sumoylation machinery, caused upregulation of NHR-25-mediated reporter expression governed by the 8xNR5RE response element. In fact, all SUMO-conjugating components tested were required for proper control of NHR-25 activity (Figs. 2 and 3). These results extend our previous findings that NHR-25 is sumoylated, and that RNAi of *smo-1* upregulates 8xNR5RE::Venus expression *in vivo*. What signals or combinations of signals normally activate or inhibit NHR-25 sumoylation are not known, nor do we know whether NHR-25 action from the 8xNR5RE synthetic response element is modulated by other signaling machineries. That is, our screen is likely far from saturation, and extending it might uncover at least some of this information.

Our recovery of the *lir-2* mutant, determination that LIR-2 is an NHR-25 co-regulator, and the biochemical and genetic demonstration that LIR-2 interacts with NHR-25 through LxxLL motifs comprise the first analyses of *lir-2* function. These studies also establish that, as in mammals and flies (see below), LxxLL is a functional NR:co-regulator interface in nematodes. The *lir-2* gene encodes a LIN-26-like (<30% identity) C2H2 zinc-finger protein (Dufourcq *et al.* 1999) in a family with two other LIN-26-related factors, LIR-1 and LIR-3. *lir-1* and *lin-26* reside in an operon and together regulate differentiation of non-neuronal ectodermal cells and the somatic gonadal epithelium (Boer *et al.* 1998; Dufourcq *et al.* 1999; Bosher *et al.* 1999). In contrast, *lir-3* is a modifier of polyglutamine aggregation in particular neurons and muscle cells (Sin *et al.* 2017). Thus, LIR proteins share similar zinc-fingers at their C-termini, but the roles of those domains are unknown, and the proteins themselves appear to be functionally distinct.

It is well established in other metazoans that co-regulators carrying LxxLL motifs, or “NR boxes”, activate or repress transcription (Heery *et al.* 1997; Nolte *et al.* 1998; Savkur and Burris 2004; Loinder and Söderström 2004; Plevin *et al.* 2005) by associating with the AF-2 domain near NR C-termini. The number, precise sequences and distinct activities of NR boxes differs among co-regulators. For example, the mammalian GRIP1/SRC2/Tif2 co-regulator contains three NR boxes (NRB1-3) that bind differentially to steroid receptors (Darimont *et al.* 1998; Kotaja 2002; He and Wilson 2003), glucocorticoid and progesterone receptors preferentially with NRB3, and estradiol receptor with NRB2 (He and Wilson 2003). Importantly, SF-1 and LRH-1, the mammalian orthologs of NHR-25, bind GRIP1 and multiple additional co-regulators through NR boxes (Børud *et al.* 2002; Suzuki *et al.* 2003; Sablin *et al.* 2003; Steffensen *et al.* 2004; Safi *et al.* 2005).

Mammalian NR boxes have been further sub-classified into four groups based on amino acid residues flanking the core motif (Chang *et al.* 1999; Suzuki *et al.* 2003; Savkur and Burris 2004). SF-1 and LRH-1 preferentially interact with sub-class III (Table 1; Suzuki *et al.* 2003). LIR-2 Lx1 is also a class III NR box with Ser at −2 and Thr at +6, whereas LIR-2 Lx2 is a class II NR box with Pro at −2. Thus, perhaps LIR-2 also binds another NR selectively through its class II NR box; several mammalian NRs preferentially associate with class II NR boxes (Chang *et al.* 1999; Kane and Means 2000). It will be interesting to determine whether LIR-2 co-regulates multiple *C. elegans* NRs or is highly selective for NHR-25. AF-2 domain sequences among NR5A family members are highly conserved (Fig. 8), and a mammalian LxxLL-AF-2 interface has been structurally determined (Mays *et al.* 2017), showing an NR helix 3-helix 12 charge clamp; SWISS-MODEL 3D structure prediction (Bienert *et al.*; Waterhouse *et al.*) suggests that NHR-25 may form a similar AF-2 domain (Fig. S4, Li *et al.* 2003; Gallastegui *et al.* 2015; Mays *et al.* 2017) to interact with LIR-2.

**Fig. 7.**
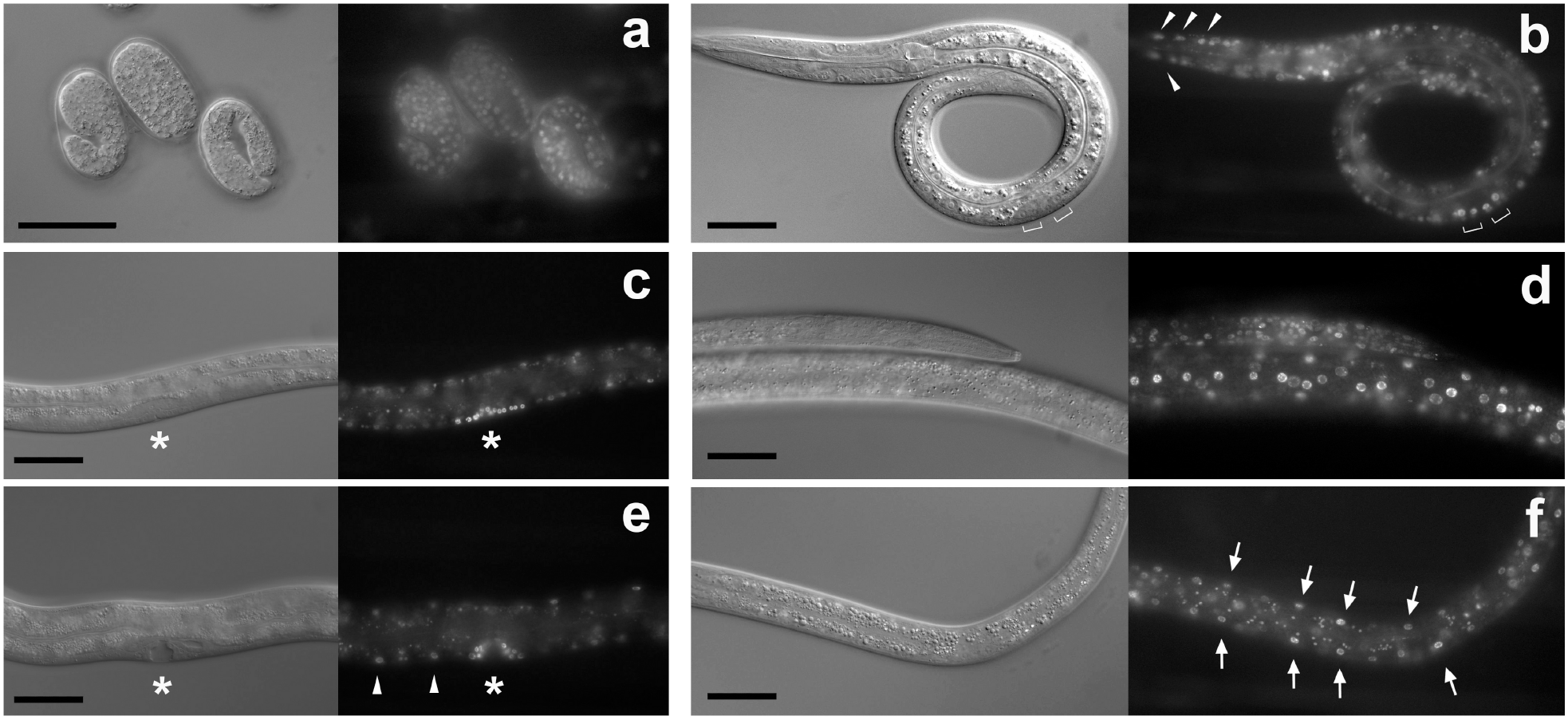
*lir-2* is expressed in overlapping tissue and cells where *nhr-25* is expressed. Expression of *lir-2a::H1-wCherry* transgene (RW11401 strain) was observed. *lir-2* is expressed ubiquitously in embryo (a), in hypodermal cells in the head (b, arrowheads), in somatic gonadal precursors (SGPs, brackets in b indicate Z1/Z4 daughters), developing vulval cells (c and e), in ventral hyp7 (e, arrowheads), and in the lateral seam cells and hyp7 cells (d) and those are all overlapping tissue and cell types where NHR-25 is expressed. In addition, *lir-2* expression was also seen in the muscle cells (f, arrows) and neurons (seen in b and d). Scale bars: 50 μm.

**Fig. 8.**
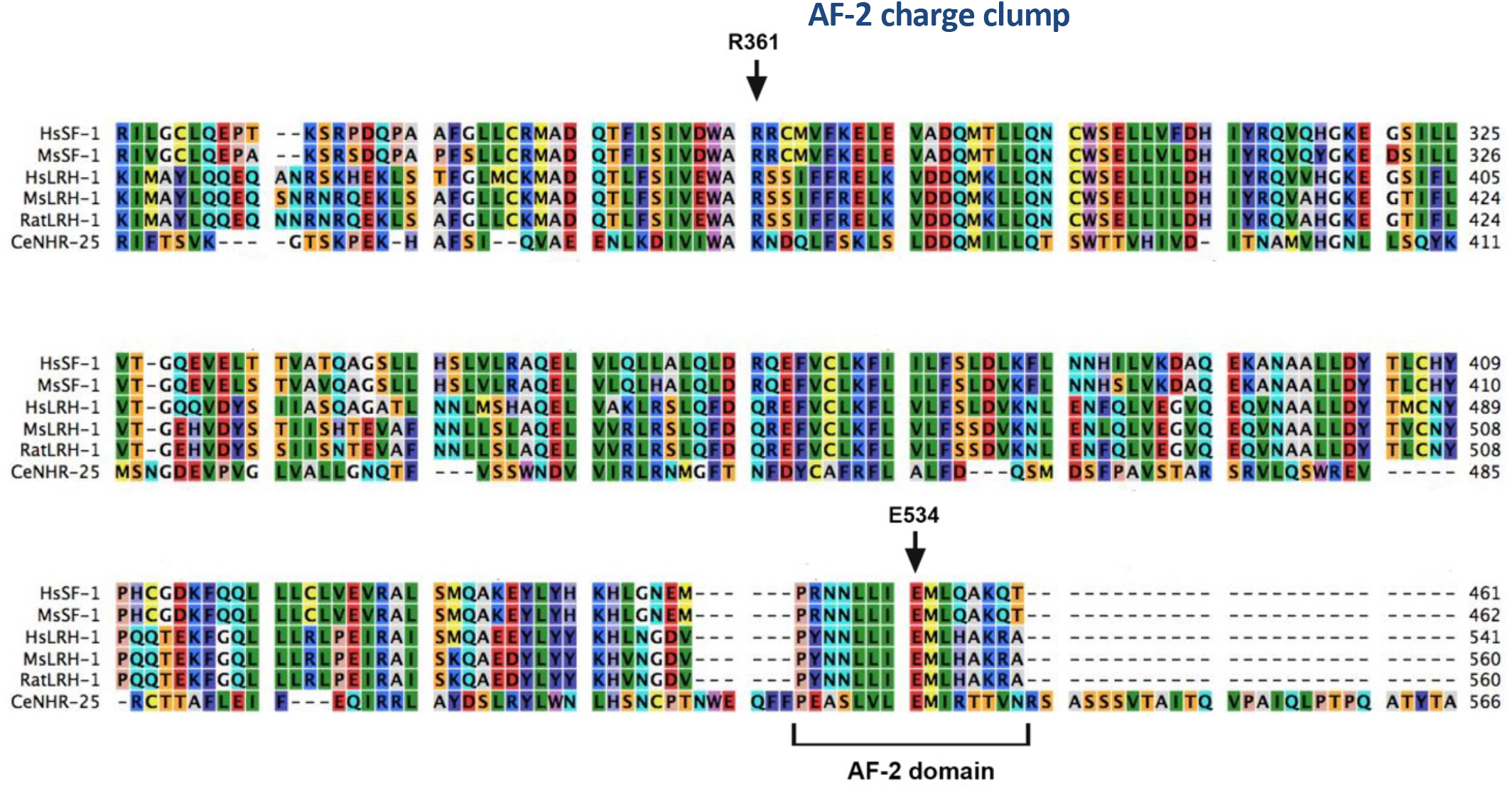
Alignment of LBD of NR5A family. Amino acid 335-566 of *C. elegans* NHR-25 was aligned with mouse (Ms) and human (Hs) SF-1 and LRH-1. R361 (in helix 3) and E534 (AF-2 domain in helix 12) in human LHR-1 were demonstrated to form the charge clamp (Mays *et al*., 2017). It is noted that most of utilizes lysine instead of arginine at R361 position as in NHR-25. Amino acid residues are colored Rasmol colors based on protein properties.

**Table 1.**
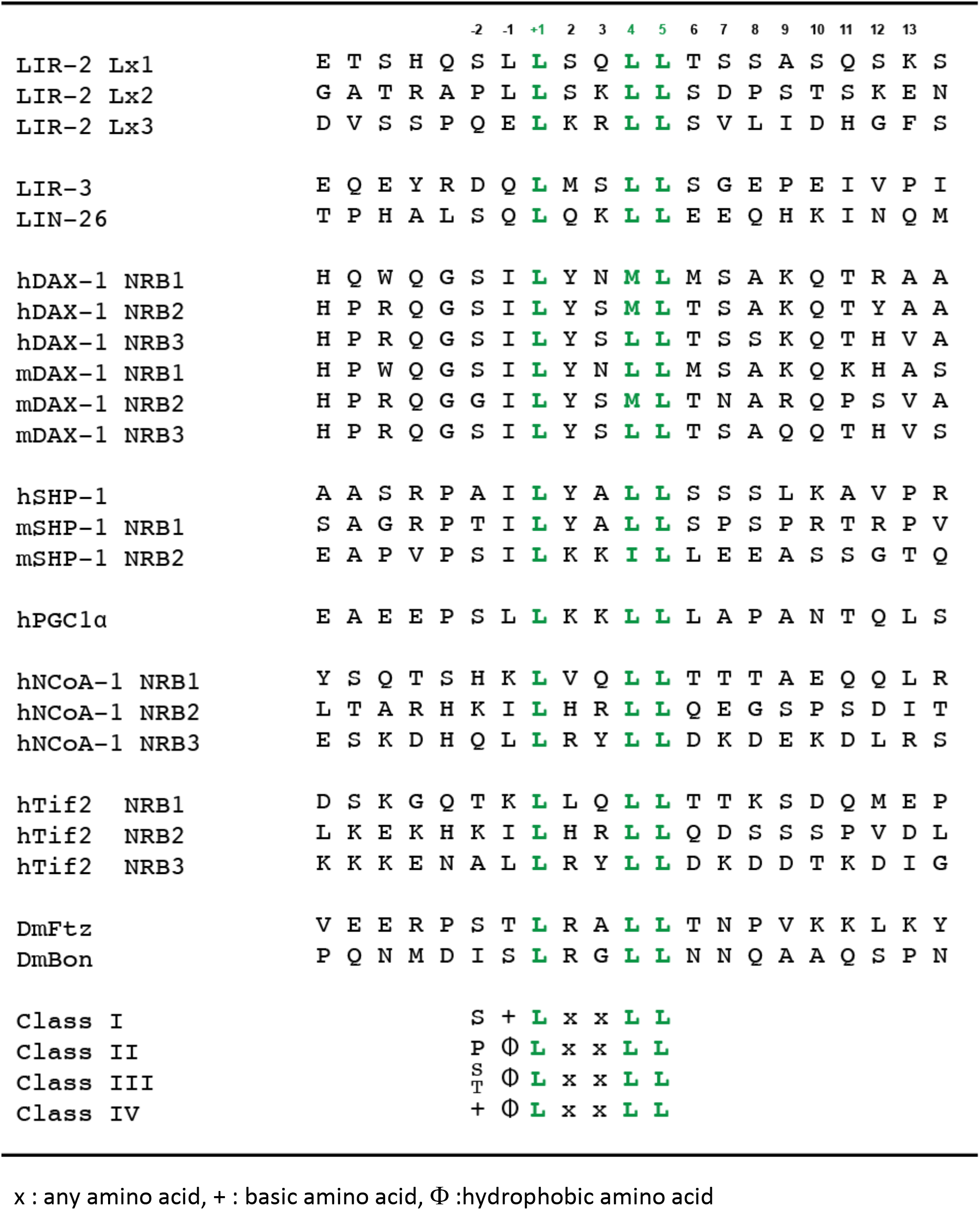
Alignment of the amino acid sequences of the NR boxes of NR5A interacting proteins

Transcription of *lir-2* is strong in early embryonic stages and decreases after the 350-cell stage, when robust expression of *lir-1/lin-26* operon transcripts and LIN-26 protein are observed (Dufourcq *et al.* 1999). Moreover, reduction of function of *lin-26* causes defects in epidermal differentiation and eventual embryonic lethality similar to observations in NHR-25 mutants (Labouesse *et al.* 1994). Given our finding that our *lir-2* mutant produced aberrant embryonic epidermal expression of the 8xNR5RE::3xVenus reporter gene, we wondered if *lir-1/lin-26* expression might be held in check by NHR-25:LIR-2 complexes in embryos and activated upon decline in *lir-2* expression. Intriguingly, ChIP-seq revealed an NHR-25 occupied region (modENCODE, D. T.-S., unpublished observations) in a highly conserved segment of the first intron of the *lir-1/lin-26* transcription unit (Fig. S1A) that includes two NR5 consensus binding sequences (Fig. S1B). However, mutants deleting one or both elements by Cas9 editing (Fig. S1C) had no apparent effect on embryogenesis or brood size (Fig. S1D). Similarly, no embryonic lethality was observed with *lir-2* mutants.

While defects either in sumoylation or in LIR-2 produced upregulation of Venus expression in similar patterns of NHR-25 expressing tissues and cell types, strikingly differences were observed in the epidermal seam cells and developing vulval cells: RNAi of any sumoylation component (*smo-1*, *uba-2*, *aos-1*, *ubc-9* and *gei-17*) enhanced Venus expression in the seam cells (Fig. 3) but not in *lir-2* mutants at any developmental stage, except for very low expression levels occasionally detected in the nuclei of dividing seam cells (Fig. 4 cr-158). Similarly, developing vulval cells (P5-7.pxx cells in mid L3 stage), which express high levels of NHR-25, lacked Venus expression in *lir-2* mutants, but showed strong upregulation of reporter expression in *smo-1* RNAi-treated animals (Fig. 4 cr178, *smo-1* RNAi panels). It is not clear the nature of these differences. As both SMO-1 and LIR-2 are expressed ubiquitously (Dufourcq *et al.* 1999; Broday 2004; Fig. 7), the mechanisms by which SUMO and LIR-2 suppress activation by NHR-25 appear to be distinct and tissue-selective.

In addition to the co-regulator and signaling enzyme identified in our genetic screen for molecules affecting NHR-25 TRC structure and function, we had shown previously that NHR-25 interacts genetically and physically with β-catenins WRM-1 and SYS-1 (Asahina *et al.* 2006) to effect cell fate decisions in the somatic gonad of *C. elegans*. In those cellular and response element contexts, WRM-1 inhibits, while SYS-1 stimulates NHR-25-dependent transcription, and NHR-25 antagonizes Wnt signaling effector POP-1/TCF-dependent transcription. Depletion of NHR-25 drives an “all-distal” cell fate in somatic gonad precursors, whereas POP-1/β-catenin loss of function mutants produce an “all-proximal” symmetrical-sister (Sys) phenotype. The NHR-25 LBD is essential for its interaction with SYS-1 and for inhibition of SYS-1-dependent activation of a POP-1/TCF reporter target gene (Asahina *et al.* 2006). Interestingly, a crystal structure of β-catenin-bound human LRH-1 (Yumoto *et al.* 2012) showed that the interaction surface is distinct from the AF-2 domain employed by LxxLL-containing co-regulators. SUMO protein interaction occurs in the NHR-25 hinge region and possibly the DNA binding domain (Ward *et al.* 2013). Thus, NHR-25 appears to employ multiple distinct surfaces, simultaneously or in discrete combinations, to receive and integrate signals and nucleate context-dependent TRC assembly.

More generally, these findings suggest that it should be possible in future work to screen on a natural NHR-25 response element, in chromosomal context, either by tracking natural target gene expression, or by inserting a screenable reporter or a selectable marker, to uncover factors that affect NHR-25 activity either positively and negatively, including components of cognate TRCs and of relevant signaling pathways. Once these are in hand, comparisons can be made to define differences and similarities in the response element, TRC factors and signaling for the same gene in different context, or in different genes in the same context. Notably, this approach is generalizable to other TFs and biological systems, including normal and disease-affected human genes, where identification of affected components could profoundly affect therapeutic development.

## ACKNOWLEDGEMENTS

We thank Kaveh Ashrafi for his generous support for microscope facility and advice. We appreciate Kirk Ehmsen and Matthew Knuesel for sharing the reagents and technical advice for CRISPR-Cas9 editing, MiSeq and immunoprecipitation; Jordan Ward for advice for CRISPR-Cas9 genome editing; Teresita Bernal and Soledad DeGuzman for preparing reagents necessary for this study. The work was supported by a grant from the National Science Foundation (MCB-1615826) to KRY. Some C. elegans strains were provided by the CGC, which is funded by NIH Office of Research Infrastructure Programs (P40 OD010440).

---

**Figure S1.**
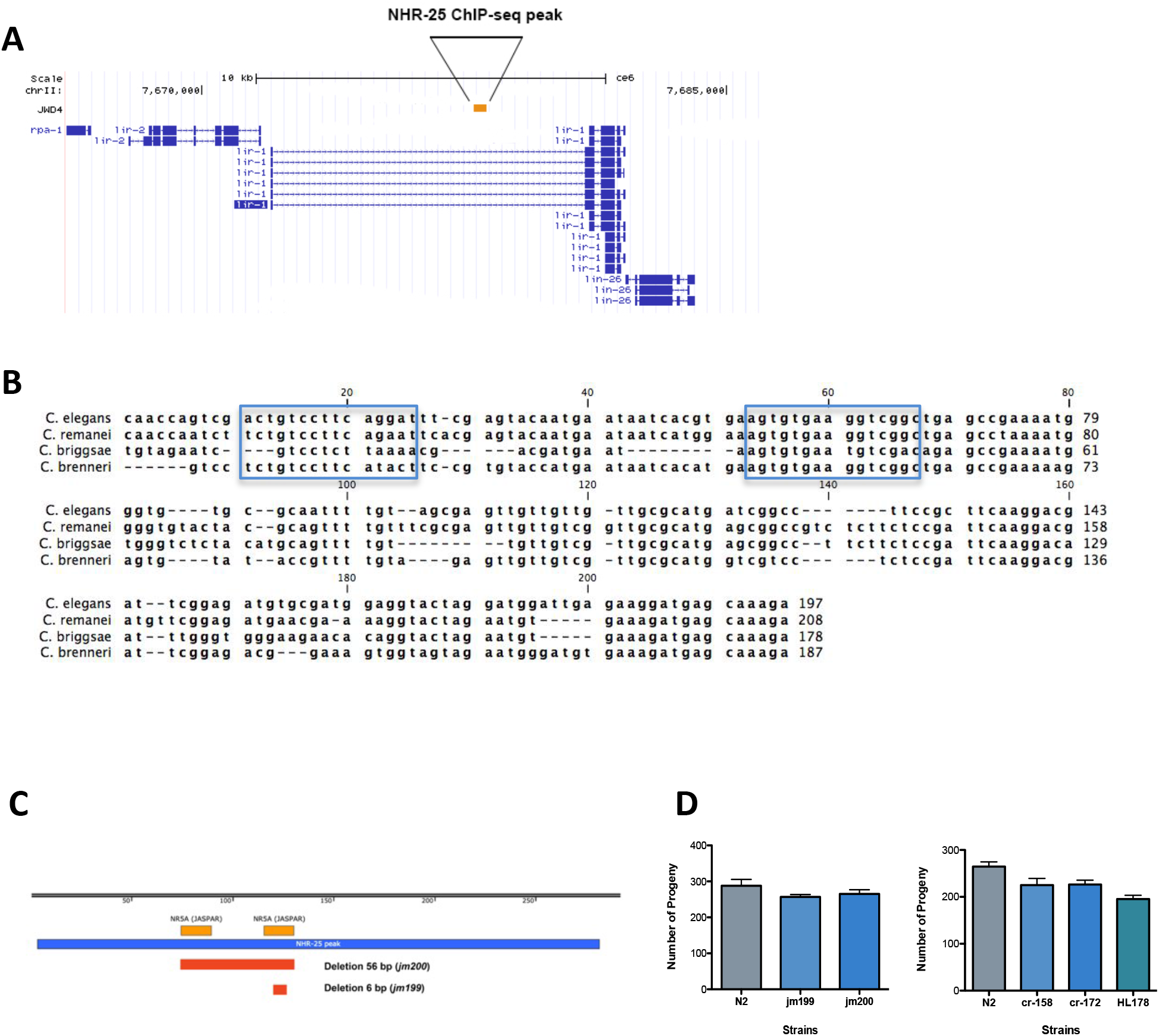
Schematic representation of upstream region of *lir-1*/*lin-26* structure. (A) *lir-1* and *lin-26* genes are located downstream of *lir-2*. NHR-25 ChIP-seq data (Thurtle-Schmidt et al., unpublished data) revealed NHR-25 occupied peak found in the long intron of *lir-1* at about 3 kb upstream of the *lir-1*/*lin-26* operon. (B) The NHR-25 ChIP-seq peak sequence is highly conserved with other *Caenorhabditis* species (Multiz alignment, UCSC Genome Browser) and two NR5A-recognition consensus are found (boxed in blue) by JASPAR2020 (scan against NR5A2 matrix ID MA0505.1). (C) depicts NHR-25 ChIP-seq peak (blue bar), NR5A recognition elements (NR5ARE) are shown (light orange bars). Dark orange bars indicate legions in CRISPR-Cas9 generated deletion mutants (jm199, jm200 strains). (D) Brood size analyses of NR5ARE deletion mutant strains (left), CRISPR-Cas9 generated *lir-2* mutants and original strain HL178 (right) at 20°C. 5-8 animals for each strain were scored. Standard error of the mean (SEM) is shown for each bar.

**Figure S2.**
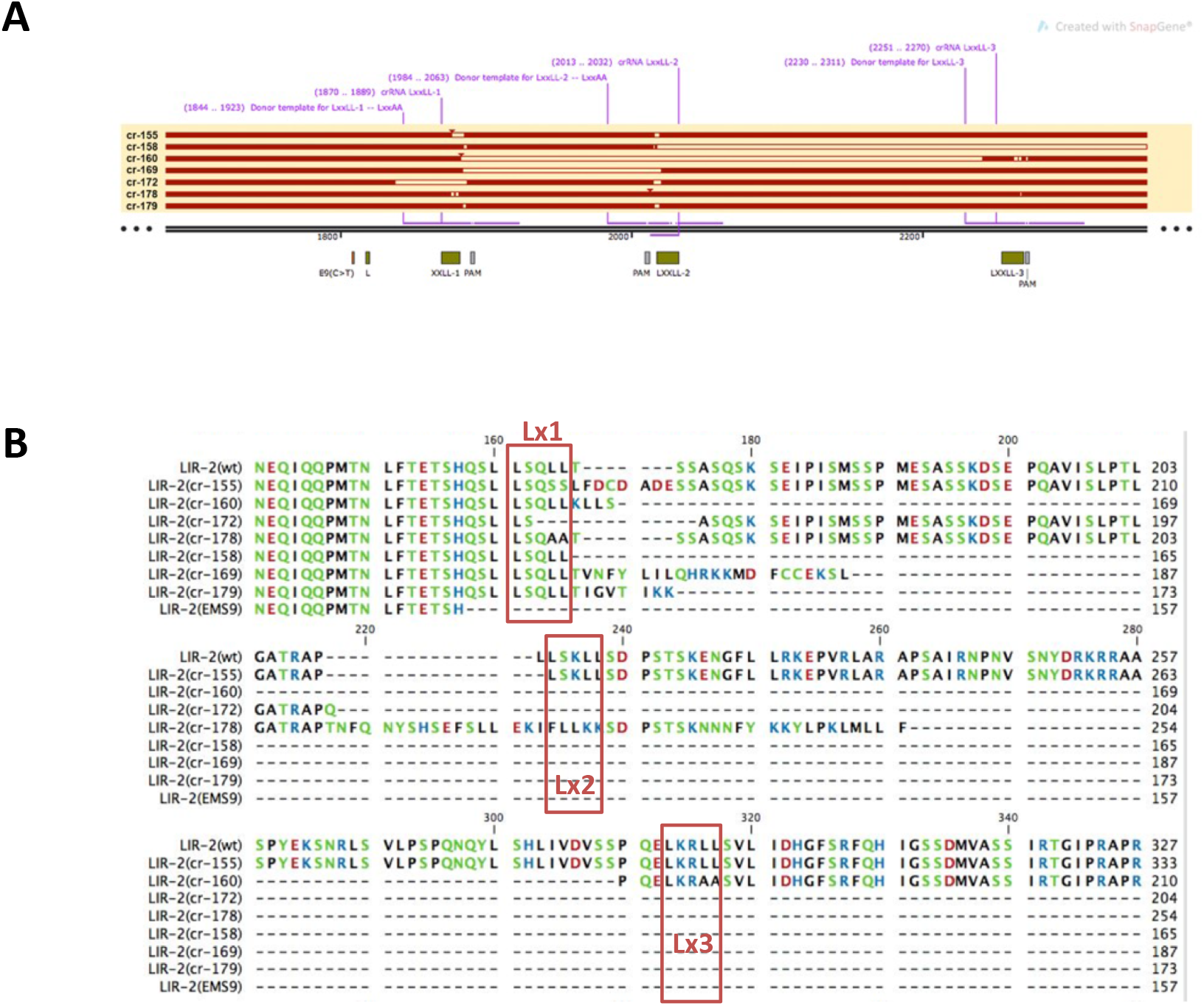
CRISPR-generated *lir-2* mutants. (A) The mutations in DNA structure in each strain are shown. crRNA and ssODN location for each LxxLL is indicated above (purple). The location of LxxLL motifs and PAM sequences are indicated below the bar graph. (B) The amino acid changes in each *lir-2* mutants are shown. Only the region containing the LxxLL motifs are shown here.

**Figure S3.**
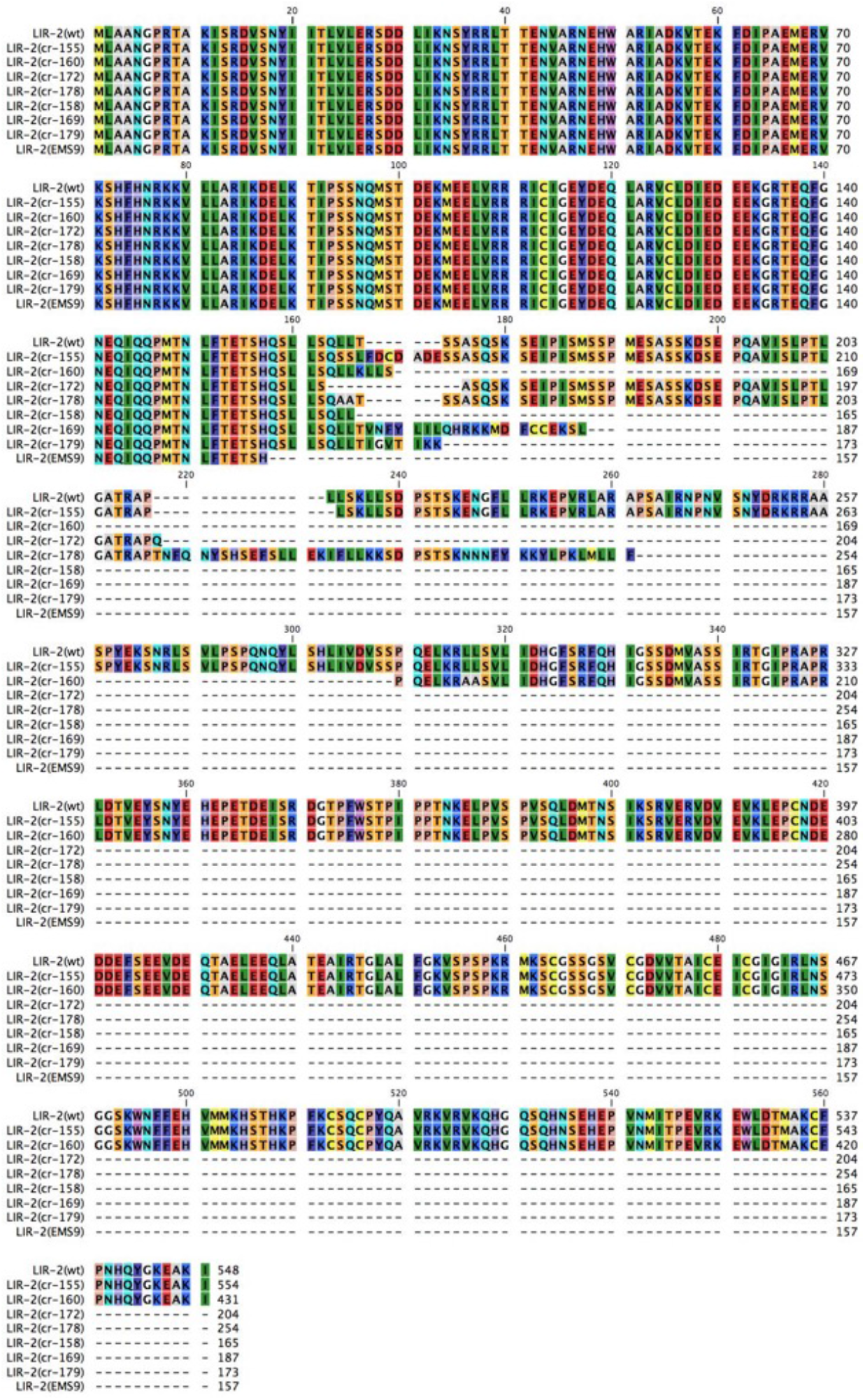
Protein alignments of CRISPR-generated *lir-2* mutants. Entire protein sequences for newly generated *lir-2* mutants are shown. Amino acid residues are colored with Rasmol colors based on protein properties.

**Figure S4.**
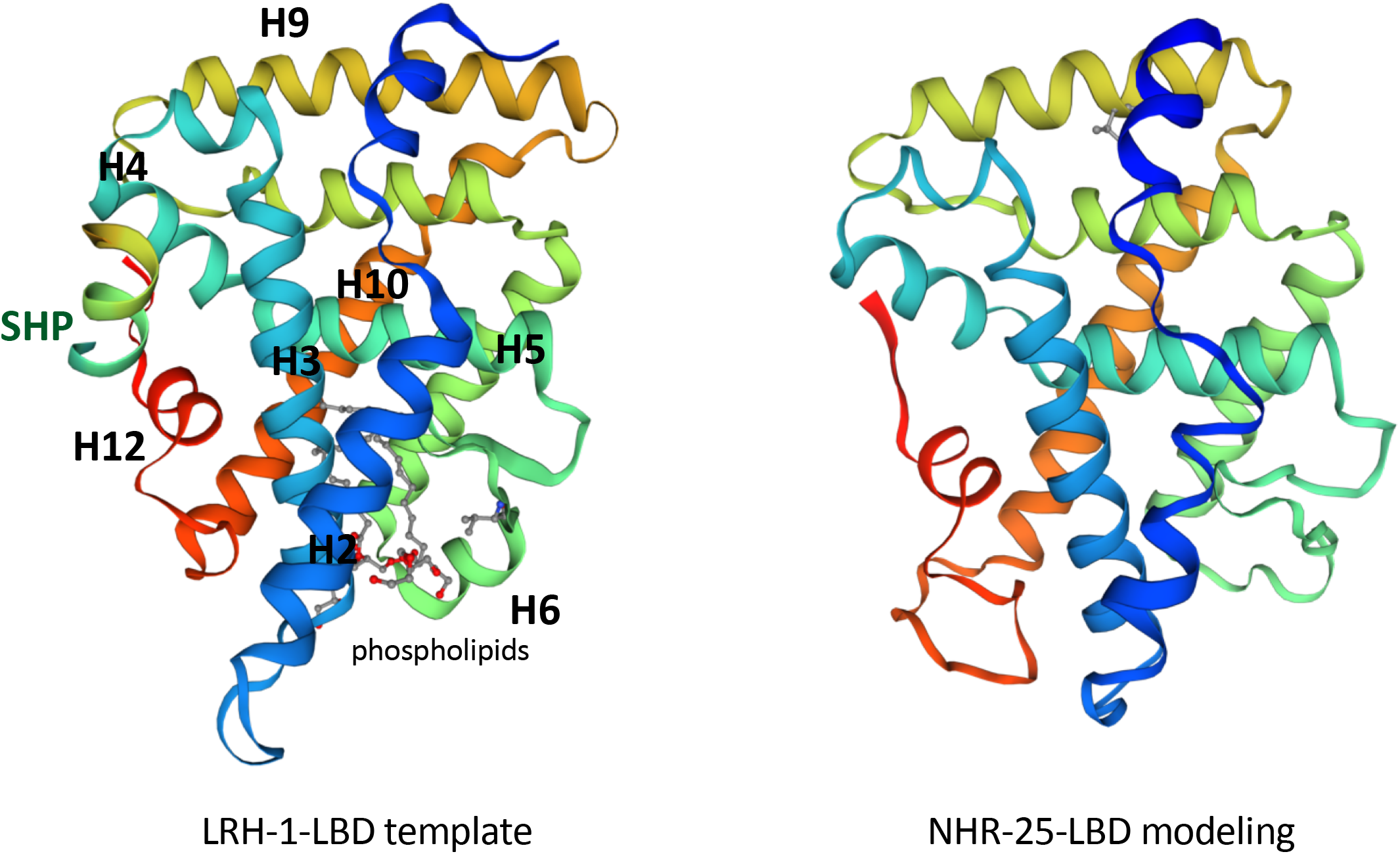
SWISS-MODEL 3D protein structure prediction for NHR-25. 3D protein model for NHR-25 was generated by homology modelling (SWISS-MODEL repository, BIOZENTRUM). It predicted that NHR-25 LBD can fold similar to that of LRH-1. LRH-1-LBD template shows corepressor SHP (small heterodimer partner 1) and phospholipids binding pocket as indicated. Helix numbers are also indicated (H2-H12).

**Table S1.**
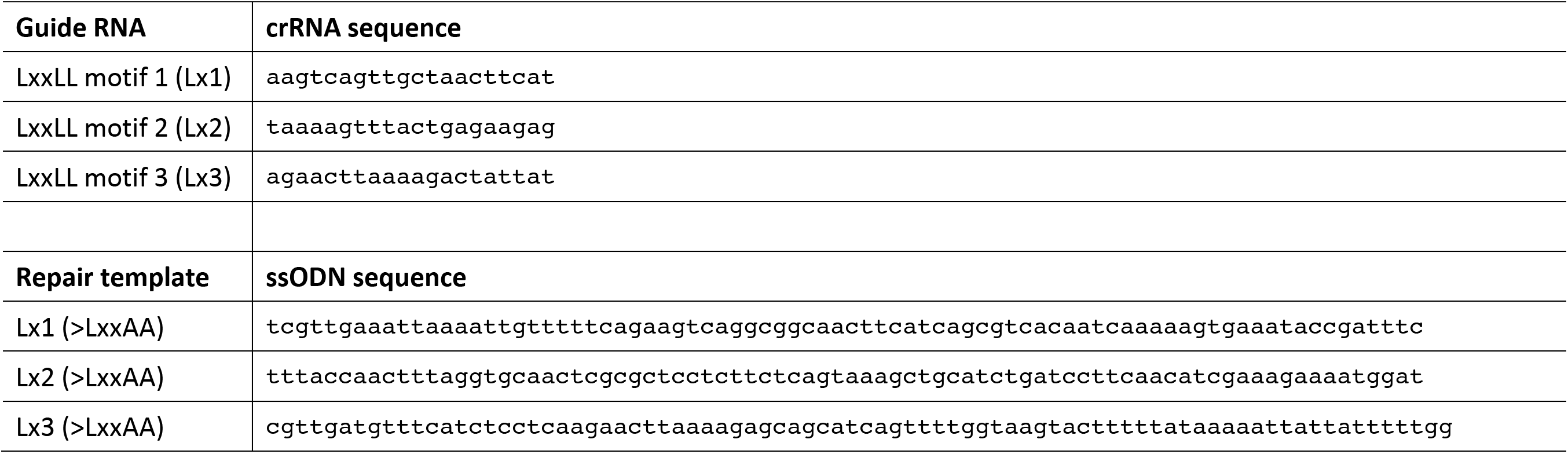
crRNA and ssODN sequences used for CRISPR-Cas9 editing in this study

**Table S2.** Number of sequencing reads mapped

| Strain | Number of Aligned Reads |
| --- | --- |
| HL178 | 267428508 |
| E2 | 216155013 |
| E9 | 159126006 |

## SUPPLEMENTAL MATERIALS

### METHODS

#### Brood size

L4 hermaphrodites were singly placed onto 35 mm NGM plates at 20 °C. Each worm was transferred to a fresh plate every 12 hr until egg laying stopped. After 24 hr of each transfer, the number of viable worms and dead embryos were scored. 5-8 worms were scored for each strain.

## REFERENCES

Antebi. A., 2015 Nuclear receptor signal transduction in C. elegans. Wormbook: 1–49.

Asahina, M., T. Ishihara, M. Jindra, Y. Kohara, I. Katsura et al., 2000 The conserved nuclear receptor Ftz-F1 is required for embryogenesis, moulting and reproduction in Caenorhabditis elegans. Genes Cells 5: 711–723.

Asahina, M., T. Valenta, M. Silhankova, V. Korinek, and M. Jindra, 2006 Crosstalk between a nuclear receptor and beta-catenin signaling decides cell fates in the C. elegans somatic gonad. Dev. Cell 11: 203–211.

Berrabah, W., P. Aumercier, P. Lefebvre, and B. Staels, 2011 Control of nuclear receptor activities in metabolism by post-translational modifications. FEBS Letters 585: 1640–1650.

Bienert, S., Waterhouse, A., de Beer, T.A.P., Tauriello, G., Studer, G., Bordoli, L., Schwede, T., 2017 The SWISS-MODEL Repository – new features and functionality. Nucleic Acids Res. 45, D313–D319.

Boer, den, B. G., S. Sookhareea, P. Dufourcq, and M. Labouesse, 1998 A tissue-specific knock-out strategy reveals that lin-26 is required for the formation of the somatic gonad epithelium in Caenorhabditis elegans. Development 125: 3213–3224.

Bosher, J. M., P. Dufourcq, S. Sookhareea, and M. Labouesse, 1999 RNA interference can target pre-mRNA: consequences for gene expression in a Caenorhabditis elegans operon. Genetics 153: 1245–1256.

Brenner, S., 1974 The genetics of Caenorhabditis elegans. Genetics 77: 71–94.

Broday, L., 2004 The small ubiquitin-like modifier (SUMO) is required for gonadal and uterine-vulval morphogenesis in Caenorhabditis elegans. Gene Dev 18: 2380–2391.

Budi, E. H., D. Duan, and R. Derynck, 2017 Transforming Growth Factor-β Receptors and Smads: Regulatory Complexity and Functional Versatility. Trends Cell Biol. 27: 658–672.

Børud, B., T. Hoang, M. Bakke, A. L. Jacob, J. Lund et al., 2002 The nuclear receptor coactivators p300/CBP/cointegrator-associated protein (p/CIP) and transcription intermediary factor 2 (TIF2) differentially regulate PKA-stimulated transcriptional activity of steroidogenic factor 1. Molecular Endocrinology 16: 757–773.

Chang, C. Y., J. D. Norris, H. Grøn, L. A. Paige, P. T. Hamilton et al., 1999 Dissection of the LXXLL nuclear receptor-coactivator interaction motif using combinatorial peptide libraries: discovery of peptide antagonists of estrogen receptors alpha and beta. Mol. Cell. Biol. 19: 8226–8239.

Chen, Z., D. J. Eastburn, and M. Han, 2004 The Caenorhabditis elegans nuclear receptor gene nhr-25 regulates epidermal cell development. Mol. Cell. Biol. 24: 7345–7358.

Darimont, B. D., R. L. Wagner, J. W. Apriletti, M. R. Stallcup, P. J. Kushner et al., 1998 Structure and specificity of nuclear receptor-coactivator interactions. Gene Dev 12: 3343–3356.

Dokshin, G. A., K. S. Ghanta, K. M. Piscopo, and C. C. Mello, 2018 Robust Genome Editing with Short Single-Stranded and Long, Partially Single-Stranded DNA Donors in Caenorhabditis elegans. Genetics 210: 781–787.

Dufourcq, P., P. Chanal, S. Vicaire, E. Camut, S. Quintin et al., 1999 lir-2, lir-1 and lin-26 encode a new class of zinc-finger proteins and are organized in two overlapping operons both in Caenorhabditis elegans and in Caenorhabditis briggsae. Genetics 152: 221–235.

Ehmsen, K. T., M. T. Knuesel, D. Martinez, M. Asahina, H. Aridomi et al., 2019 Definition of alleles and altered regulatory motifs across Cas9-edited cell populations. bioRxiv 7: 146–48.

Gallastegui, N., J. A. G. Mackinnon, R. J. Fletterick, and E. Estébanez-Perpiñá, 2015 Advances in our structural understanding of orphan nuclear receptors. Trends in Biochemical Sciences 40: 25–35.

Gissendanner, C. R., and A. E. Sluder, 2000 nhr-25, the Caenorhabditis elegans ortholog of ftz-f1, is required for epidermal and somatic gonad development. Dev. Biol. 221: 259–272.

Hada, K., M. Asahina, H. Hasegawa, Y. Kanaho, F. J. Slack et al., 2010 The nuclear receptor gene nhr-25 plays multiple roles in the Caenorhabditis elegans heterochronic gene network to control the larva-to-adult transition. Dev. Biol. 344: 1100–1109.

Hajduskova, M., M. Jindra, M. A. Herman, and M. Asahina, 2009 The nuclear receptor NHR-25 cooperates with the Wnt/beta-catenin asymmetry pathway to control differentiation of the T seam cell in C. elegans. J. Cell. Sci. 122: 3051–3060.

Hayes, G. D., A. R. Frand, and G. Ruvkun, 2006 The mir-84 and let-7 paralogous microRNA genes of Caenorhabditis elegans direct the cessation of molting via the conserved nuclear hormone receptors NHR-23 and NHR-25. Development 133: 4631–4641.

He, B., and E. M. Wilson, 2003 Electrostatic modulation in steroid receptor recruitment of LXXLL and FXXLF motifs. Mol. Cell. Biol. 23: 2135–2150.

Heery, D. M., E. Kalkhoven, S. Hoare, and M. G. Parker, 1997 A signature motif in transcriptional co-activators mediates binding to nuclear receptors. Nature 387: 733–736.

Kamath, R. S., A. G. Fraser, Y. Dong, G. Poulin, R. Durbin et al., 2003 Systematic functional analysis of the Caenorhabditis elegans genome using RNAi. Nature 421: 231–237.

Kane, C. D., and A. R. Means, 2000 Activation of orphan receptor-mediated transcription by Ca(2+)/calmodulin-dependent protein kinase IV. EMBO J. 19: 691–701.

Kastner, P., M. Mark, and P. Chambon, 1995 Nonsteroid nuclear receptors: what are genetic studies telling us about their role in real life? Cell.

Kotaja, N., 2002 The Nuclear Receptor Interaction Domain of GRIP1 Is Modulated by Covalent Attachment of SUMO-1. Journal of Biological Chemistry 277: 30283–30288.

Kramer, J. M., R. P. French, E. C. Park, and J. J. Johnson, 1990 The Caenorhabditis elegans rol-6 gene, which interacts with the sqt-1 collagen gene to determine organismal morphology, encodes a collagen. Mol. Cell. Biol. 10: 2081–2089.

Labouesse, M., S. Sookhareea, and H. R. Horvitz, 1994 The Caenorhabditis elegans gene lin-26 is required to specify the fates of hypodermal cells and encodes a presumptive zinc-finger transcription factor. Development 120: 2359–2368.

Lee, J. H., and M. J. Lee, 2012 Emerging roles of the ubiquitin-proteasome system in the steroid receptor signaling. Arch. Pharm. Res. 35: 397–407.

Li, H., and R. Durbin, 2009 Fast and accurate short read alignment with Burrows-Wheeler transform. Bioinformatics 25: 1754–1760.

Li, Y., M. H. Lambert, and H. E. Xu, 2003 Activation of Nuclear Receptors: A Perspective from Structural Genomics. Structure 11: 741–746.

Loinder, K., and M. Söderström, 2004 Functional analyses of an LXXLL motif in nuclear receptor corepressor (N-CoR). J. Steroid Biochem. Mol. Biol. 91: 191–196.

Lois, L. M., and C. D. Lima, 2005 Structures of the SUMO E1 provide mechanistic insights into SUMO activation and E2 recruitment to E1. EMBO J. 24: 439–451.

Lonard, D. M., and B. W. O’Malley, 2012 Nuclear receptor coregulators: modulators of pathology and therapeutic targets. Nat Rev Endocrinol 8: 598–604.

Mangelsdorf, D. J., C. Thummel, M. Beato, P. Herrlich, G. Schütz et al., 1995 The nuclear receptor superfamily: the second decade. Cell 83: 835–839.

Mays, S. G., C. D. Okafor, M. L. Tuntland, R. J. Whitby, V. Dharmarajan et al., 2017 Structure and Dynamics of the Liver Receptor Homolog 1-PGC1α Complex. Mol. Pharmacol. 92: 1–11.

McInerney, E. M., D. W. Rose, S. E. Flynn, S. Westin, T. M. Mullen et al., 1998 Determinants of coactivator LXXLL motif specificity in nuclear receptor transcriptional activation. Gene Dev 12: 3357–3368.

McNally, J. G., W. G. Müller, D. Walker, R. Wolford, and G. L. Hager, 2000 The Glucocorticoid Receptor: Rapid Exchange with Regulatory Sites in Living Cells. Science 287: 1262–1265.

Mello, C., and A. Fire, 1995 DNA transformation. Methods in cell biology Caenorhabditis elegans: modern biological analysis of an organism 48: 451–482.

Monsalve, G. C., and A. R. Frand, 2012 Toward a unified model of developmental timing: A “molting” approach. worm 1: 221–230.

Mullaney, B. C., R. D. Blind, G. A. Lemieux, C. L. Pérez, I. C. Elle et al., 2010 Regulation of C. elegans Fat Uptake and Storage by Acyl-CoA Synthase-3 Is Dependent on NR5A Family Nuclear Hormone Receptor nhr-25. Cell Metab. 12: 398–410.

Nelson, M. D., E. Zhou, K. Kiontke, H. Fradin, G. Maldonado et al., 2011 A bow-tie genetic architecture for morphogenesis suggested by a genome-wide RNAi screen in Caenorhabditis elegans. PLoS Genet. 7: e1002010.

Nolte, R. T., G. B. Wisely, S. Westin, J. E. Cobb, M. H. Lambert et al., 1998 Ligand binding and co-activator assembly of the peroxisome proliferator-activated receptor-gamma. Nature 395: 137–143.

Paix, A., A. Folkmann, D. Rasoloson, and G. Seydoux, 2015 High Efficiency, Homology-Directed Genome Editing in Caenorhabditis elegans Using CRISPR-Cas9 Ribonucleoprotein Complexes. Genetics 201: 47–54.

Plevin, M. J., M. M. Mills, and M. Ikura, 2005 The LxxLL motif: a multifunctional binding sequence in transcriptional regulation. Trends in Biochemical Sciences 30: 66–69.

Rual, J.-F., J. Ceron, J. Koreth, T. Hao, A.-S. Nicot et al., 2004 Toward improving Caenorhabditis elegans phenome mapping with an ORFeome-based RNAi library. Genome Res. 14: 2162–2168.

Sablin, E. P., I. N. Krylova, R. J. Fletterick, and H. A. Ingraham, 2003 Structural basis for ligand-independent activation of the orphan nuclear receptor LRH-1. Molecular Cell 11: 1575–1585.

Safi, R., A. Kovacic, S. Gaillard, Y. Murata, E. R. Simpson et al., 2005 Coactivation of liver receptor homologue-1 by peroxisome proliferator-activated receptor gamma coactivator-1alpha on aromatase promoter II and its inhibition by activated retinoid X receptor suggest a novel target for breast-specific antiestrogen therapy. Cancer Research 65: 11762–11770.

Savkur, R. S., and T. P. Burris, 2004 The coactivator LXXLL nuclear receptor recognition motif. J. Pept. Res. 63: 207–212.

Silhankova, M., M. Jindra, and M. Asahina, 2005 Nuclear receptor NHR-25 is required for cell-shape dynamics during epidermal differentiation in Caenorhabditis elegans. J. Cell. Sci. 118: 223–232.

Sin, O., T. de Jong, A. Mata-Cabana, M. Kudron, M. A. Zaini et al., 2017 Identification of an RNA Polymerase III Regulator Linked to Disease-Associated Protein Aggregation. Molecular Cell 65: 1096–1108.e6.

Steffensen, K. R., E. Holter, A. Båvner, M. Nilsson, M. Pelto-Huikko et al., 2004 Functional conservation of interactions between a homeodomain cofactor and a mammalian FTZ-F1 homologue. EMBO Rep. 5: 613–619.

Suzuki, T., M. Kasahara, H. Yoshioka, K.-I. Morohashi, and K. Umesono, 2003 LXXLL-related motifs in Dax-1 have target specificity for the orphan nuclear receptors Ad4BP/SF-1 and LRH-1. Mol. Cell. Biol. 23: 238–249.

Timmons, L., D. L. Court, and A. Fire, 2001 Ingestion of bacterially expressed dsRNAs can produce specific and potent genetic interference in Caenorhabditis elegans. Gene 263: 103–112.

Treuter, E., and N. Venteclef, 2011 Transcriptional control of metabolic and inflammatory pathways by nuclear receptor SUMOylation. Biochimica et Biophysica Acta (BBA) – Molecular Basis of Disease 1812: 909–918.

Wang, K., M. Li, and H. Hakonarson, 2010 ANNOVAR: functional annotation of genetic variants from high-throughput sequencing data. Nucleic Acids Research 38: e164–e164.

Ward, J. D., N. Bojanala, T. Bernal, K. Ashrafi, M. Asahina et al., 2013 Sumoylated NHR-25/NR5A Regulates Cell Fate duringC. elegans Vulval Development. PLoS Genet. 9: e1003992.

Ward, J. D., B. Mullaney, B. J. Schiller, L. D. He, S. E. Petnic et al., 2014 Defects in the C. elegans acyl-CoA synthase, acs-3, and nuclear hormone receptor, nhr-25, cause sensitivity to distinct, but overlapping stresses. PLoS ONE 9: e92552.

Waterhouse, A., Bertoni, M., Bienert, S., Studer, G., Tauriello, G., Gumienny, R., Heer, F.T., de Beer, T.A.P., Rempfer, C., Bordoli, L., Lepore, R., Schwede, T., 2018 SWISS-MODEL: homology modelling of protein structures and complexes. Nucleic Acids Res. 46, W296–W303.

Watson, L. C., K. M. Kuchenbecker, B. J. Schiller, J. D. Gross, M. A. Pufall et al., 2013 The glucocorticoid receptor dimer interface allosterically transmits sequence-specific DNA signals. Nat. Struct. Mol. Biol. 20: 876–883.

Weikum, E. R., M. T. Knuesel, E. A. Ortlund, and K. R. Yamamoto, 2017 Glucocorticoid receptor control of transcription: precision and plasticity via allostery. Nat. Rev. Mol. Cell Biol. 18: 159–174.

Yamamoto, K. R., 1985 Steroid receptor regulated transcription of specific genes and gene networks. Annu. Rev. Genet. 19: 209–252.

Yoshinaga, S. K., C. L. Peterson, I. Herskowitz, and K. R. Yamamoto, 1992 Roles of SWI1, SWI2, and SWI3 proteins for transcriptional enhancement by steroid receptors. Science 258: 1598–1604.

Yumoto, F., P. Nguyen, E. P. Sablin, J. D. Baxter, P. Webb et al., 2012 Structural basis of coactivation of liver receptor homolog-1 by β-catenin. Proceedings of the National Academy of Sciences 109: 143–148.

